# OLIG2 Drives Abnormal Neurodevelopmental Phenotypes in Human iPSC-Based Organoid and Chimeric Mouse Models of Down Syndrome

**DOI:** 10.1101/462739

**Authors:** Ranjie Xu, Andrew T Brawner, Shenglan Li, JingJing Liu, Hyosung Kim, Haipeng Xue, Zhiping P. Pang, Woo-Yang Kim, Ronald P. Hart, Ying Liu, Peng Jiang

## Abstract

Down syndrome (DS) is a common neurodevelopmental disorder, and cognitive defects in DS patients may arise form imbalances in excitatory and inhibitory neurotransmission. Understanding the mechanisms underlying such imbalances may provide opportunities for therapeutic intervention. Here, we show that human induced pluripotent stem cells (hiPSCs) derived from DS patients overproduce OLIG2^+^ ventral forebrain neural progenitors. As a result, DS hiPSC-derived cerebral organoids excessively produce specific subclasses of GABAergic interneurons and cause impaired recognition memory in neuronal chimeric mice. Increased OLIG2 expression in DS cells directly upregulates interneuron lineage-determining transcription factors. shRNA-mediated knockdown of *OLIG2* largely reverses abnormal gene expression in early-stage DS neural progenitors, reduces interneuron production in DS organoids and chimeric mouse brains, and improves behavioral deficits in DS chimeric mice. Thus, altered OLIG2 expression may underlie neurodevelopmental abnormalities and cognitive defects in DS patients.

## INTRODUCTION

Down syndrome (DS), caused by human chromosome 21 (HSA21) trisomy, is the leading genetic cause of intellectual disability (Parker et al., 2010). An imbalance in excitatory and inhibitory neurotransmission is one of the underlying causes of cognitive deficit of DS (Fernandez et al., 2007; Haydar and Reeves, 2012; Rissman and Mobley, 2011). The inhibitory GABAergic interneurons in the cerebral cortex are derived from the neuroepithelium of the embryonic ventral forebrain (Butt et al., 2005; Kessaris et al., 2006; Marin, 2012; Wonders et al., 2008). Many of these neuroepithelial cells express the HSA21 genes *OLIG1* and *OLIG2*. In mice, both Olig1 and Olig2 are expressed in the embryonic neuroepithelium of the ventral forebrain (Lu et al., 2000; Petryniak et al., 2007). In humans, OLIG2, but not OLIG1, is abundantly expressed in the embryonic ventral forebrain (Jakovcevski and Zecevic, 2005), as opposed to their overlapping expression pattern in mouse embryonic brain. Thus, establishing the role of human *OLIG* genes, in regulating interneuron production, is critical for understanding the mechanisms underlying cognitive deficit in DS and may help devise novel therapeutic strategies.

It is highly debatable how the production of GABAergic neurons is altered in DS and how *OLIG* genes are involved as regulators of GABAergic neuron production under normal and DS disease conditions. First, using mouse models, studies examining the functions of *Olig* genes in GABAergic neuron production remain inconclusive. Loss-of-function studies showed that only Olig1 repressed the production of GABAergic interneurons (Furusho et al., 2006; Ono et al., 2008; Petryniak et al., 2007; Silbereis et al., 2014). Gain-of-function studies showed that overexpression of Olig2 promoted the production of GABAergic neurons (Liu et al., 2015). However, this finding is confounded by the fact that the overexpression and mis-expression of Olig2 in inappropriate cells and developmental stages caused massive cell death in the mouse brain (Liu et al., 2015). Second, DS mouse models often show discrepancies in modeling DS-related genotype-phenotype relationships. The discrepant findings in genotype and phenotypic expression of *Olig* genes, and changes in the number of GABAergic neurons from different DS mouse models are summarized in Table S1. Third, while studies in the Ts65Dn mouse model of DS indicated that GABAergic neurons were overproduced (Chakrabarti et al., 2010) and inhibiting the GABAergic transmission could alleviate cognitive deficit (Fernandez et al., 2007), studies using postmortem brain tissues from elderly DS patients (Kobayashi et al., 1990; Ross et al., 1984) and 2-dimensional (2D) cultures of DS human induced pluripotent stem cells (hiPSCs) (Huo et al., 2018) contradictorily showed reduced production of GABAergic neurons.

The lack of availability of functional human brain tissue from normal or DS patients is preventive for a detailed mechanistic understanding of DS pathophysiology. Recent studies have demonstrated the utility of hiPSCs derived from individuals with DS as a human cellular model of DS brain development (Briggs et al., 2013; Chen et al., 2014; Jiang et al., 2013b; Shi et al., 2012; Weick et al., 2013). Moreover, the hiPSC-derived 3D brain organoids display structural organizations and cytoarchitecture resembling the developing human brain and have significantly advanced our knowledge on human brain development and pathology (Amin and Pasca, 2018; Brawner et al., 2017; Centeno et al., 2018; Simao et al., 2018). In this study, we employ brain organoid and *in vivo* chimeric mouse brain models (Chen et al., 2016) to investigate the functions of *OLIG* genes in human interneuron development and pathogenesis. Our findings suggest OLIG2 as an excellent potential target for developing personalized prenatal therapy for DS (Bianchi, 2012; de Wert et al., 2017; Guedj et al., 2014).

## RESULTS

### Human PSC-derived OLIG2^+^ ventral forebrain NPCs give rise to GABAergic neurons

To test the hypothesis that human OLIG2 is involved in interneuron development, we used OLIG2-GFP hPSC reporter lines generated in our previous studies (Liu et al., 2011; Xue et al., 2009). To obtain ventralized brain organoids, we treated organoids cultured from primitive neural progenitor cells (pNPCs) with Sonic Hedgehog (SHH) and purmorphamine (Pur), a small-molecule agonist of SHH signaling pathway, from week 3 to 5 (Figure 1A). At week 5, many cells in the organoids started to show GFP fluorescence (Figure 1A and Figure S1A). Most of the cells in the organoids expressed the NPC marker Nestin, forebrain marker FOXG1, as well as NKX2.1, a marker of ventral prosencephalic progenitors (Sussel et al., 1999; Xu et al., 2004), but not dorsal brain marker PAX6 (Ma et al., 2012) (Figures 1B, 1C), indicating that the vast majority of the cells were ventral forebrain NPCs. In these organoids, we observed distinct rosette-like structures that resemble the proliferative regions of human ventricular zone containing SOX2^+^ progenitors, and β-III-Tubulin (βIIIT^+^) immature neurons (Figure S1B). The GFP fluorescence faithfully mirrored the expression of OLIG2 in the NPCs (Figures 1B, 1C; about 38% of the total cells expressed both GFP and OLIG2). Only a very small fraction of the NPCs expressed OLIG1 (1.1 ± 0.2%, Figures 1B, 1C) and OLIG1 expression did not overlap with OLIG2/GFP and appeared to be localized in the cytoplasm (Figure 1B), which is consistent with a previous report using human fetal brain tissue (Jakovcevski and Zecevic, 2005). These observations were further confirmed by immunoblot analysis. OLIG2 was abundantly expressed and present mostly in the nuclear fraction, whereas OLIG1 was detected at a low level in the cytosolic fraction (Figure 1D). Unexpectedly, qPCR results demonstrated that during the patterning stage, the levels of both *OLIG* mRNAs were significantly increased from week 3 to 5. The discrepancy between *OLIG1* transcript level and its protein level may suggest the involvement of precise spatiotemporal control of mRNA translation observed in cortical development (Kraushar et al., 2014; Kwan et al., 2012). After initiating neuronal differentiation, *OLIG2* expression started to decrease over time and GFP fluorescence became dim at week 8 (Figure 1A and Figure S1A). Nevertheless, at 8 weeks, some of the βIIIT^+^ neurons, GABA^+^ neurons, and S100β^+^ astrocytes were labeled by GFP staining, indicating these cells were likely derived from OLIG2^+^/GFP^+^ NPCs. However, we did not find any oligodendroglial lineage cells in these organoids cultured under the neuronal differentiation condition. The expression of both *OLIG* gene transcripts significantly decreased at week 8 (Figure 1E). Organoids had been known to lack oligodendrocytes (Pasca, 2018; Quadrato et al., 2016). It was not until recently that the induction of oligodendrogenesis and myelination was achieved in organoids (Madhavan et al., 2018). In a separate set of experiments, we cultured the 5-week-old organoids under a glial differentiation condition and found MBP^+^ mature oligodendrocytes in the 8-week-old organoids (Figure S1C), demonstrating that the OLIG2^+^/GFP^+^ NPCs in the organoids had the potential to differentiate to oligodendroglial lineage cells if supplied with the appropriate signals.

**Figure 1.**
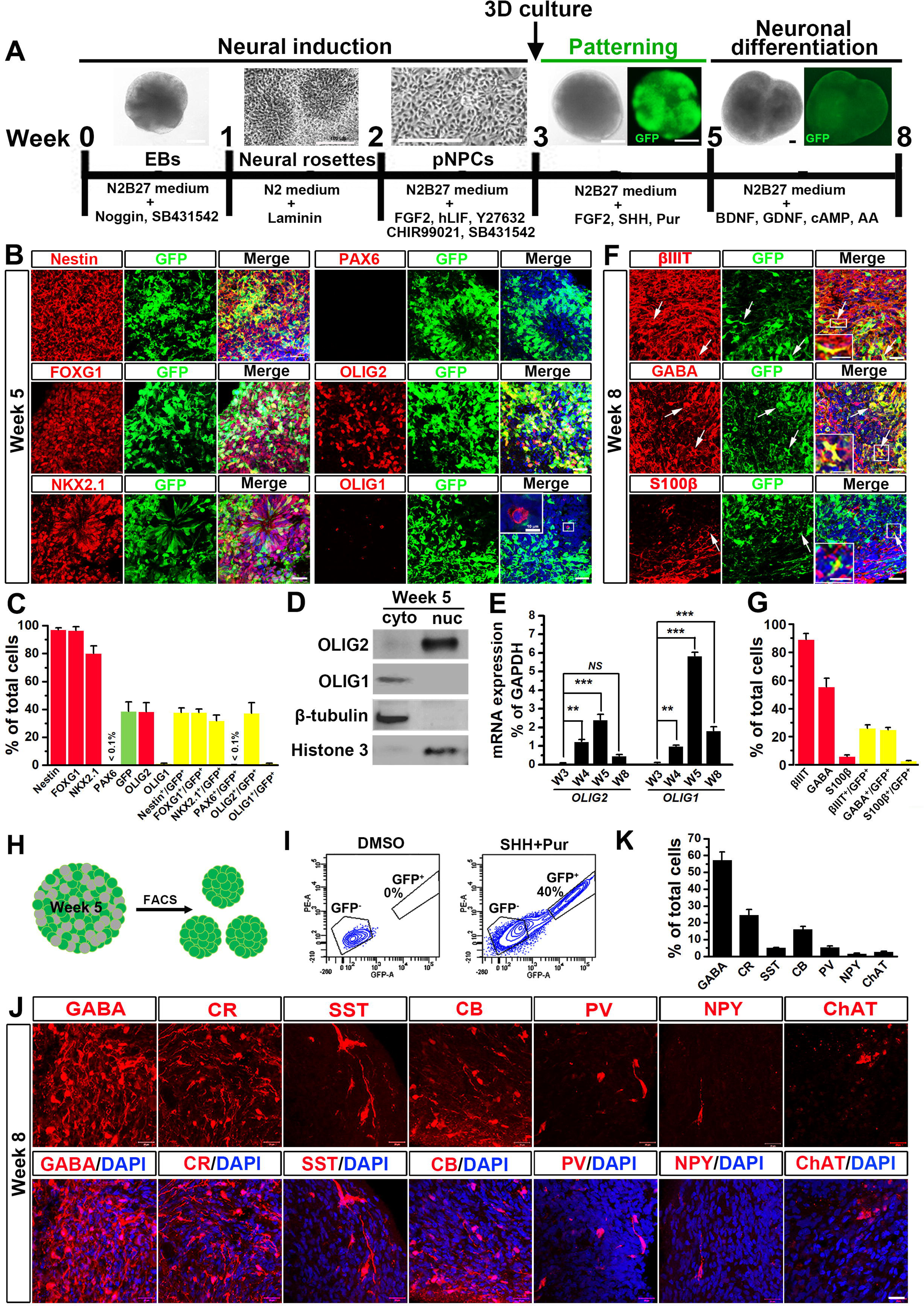
Generation of ventral forebrain organoids and GABAergic neurons from OLIG2-GFP hPSC reporter lines. (A) A schematic procedure for deriving ventral forebrain organoids by the treatment of sonic hedgehog (SHH) and purmorphamine (Pur). Scale bars: 100 μm. (B and C) Representatives and quantification of Nestin-, FOXG1-, NKX2.1-, PAX6-, OLIG2-, OLIG1- and GFP-expressing cells in 5 to 6-week-old organoids (n = 4). Scale bars: 20 μm or 10 μm in the original or enlarged images, respectively. (D) Western blotting analysis shows the expression of OLIG1 and OLIG2 in the nuclear (nuc) and cytosolic (cyto) fraction from 5-week-old organoids. (E) qPCR analysis of *OLIG2* and *OLIG1* mRNA expression in the organoids at different stages (n = 4). One-way ANOVA test. \*\**P* < 0.01, \*\*\**P* < 0.001, and *NS* represents no significance. (F and G) Representatives and quantification of βIIIT-, GABA-, S100β-, and GFP-expressing cells in 8- week-old organoids (n = 4). Scale bars: 20 μm or 10 μm in the original or enlarged images, respectively. (H) A schematic diagram showing that GFP^+^ cells are purified from 5-week-old organoids by FACS and then cultured to form organoids. (I) FACS analysis showing the percentage of GFP^+^ cells in the 5-week-old organoids treated with SHH and Pur and the control organoids treated with DMSO (n = 4). (J and K) Representatives and quantification of GABA-, CB-, CR-, PV-, SST-, NPY-, and ChAT- expressing neurons in 8-week-old GFP^+^ cell-derived organoids (n = 4). Scale bars: 20 μm. See also Figure S1.

To further delineate neuronal fate of the OLIG2^+^ NPCs, we purified the GFP^+^ cells using fluorescence-activated cell sorting (FACS). About 40% of the total cells in week 5 organoids were collected as GFP^+^ cells (Figures 1H, 1I, S1D). These cells expressed Nestin, NKX2.1, and the mid/forebrain marker OTX2, confirming that these GFP^+^ cells were OLIG2-expressing ventral telencephalic NPCs (Figure S1D). Next, we cultured the purified OLIG2^+^/GFP^+^ cells in 3D to generate organoids. GFP fluorescence was strong at week 5 but became dim at week 8 (Figure S1E). We found NeuN^+^ and S100β^+^ cells in the organoids, but not NG2^+^ or PDGFRα^+^ cells (Figures S1F, S1G). The majority of OLIG2^+^/GFP^+^ NPCs in the organoids efficiently differentiated into GABAergic neurons (Figures 1J, 1K). GABAergic interneurons are conventionally categorized based on neuropeptide and Ca^2+^ binding protein expression (Petilla Interneuron Nomenclature Group et al., 2008). We thus stained the organoids with the major subclass markers calretinin (CR), calbindin (CB), parvalbumin (PV), somatostatin (SST), and neuropeptide Y (NPY). As shown in Figures 1J, 1K, of these markers, CR was most highly expressed, and CB, SST, and PV were robustly detected. A small percentage of NPY^+^ neurons was also detected. After FACS purification, some of the OLIG2^+^/GFP^+^ NPCs were plated onto coverslips and cultured under 2D conditions in neuronal differentiation medium for 3 weeks. As opposed to 3D cultures, in the 2D cultures maintained for the same time period, the OLIG2^+^/GFP^+^ NPCs only differentiated into CR^+^ neurons, with other subclasses of GABAergic neuron not detected (Figures S1H, S1I), suggesting that the 3D organoid models might be more effective in recapitulating a broader range of GABAergic neuron differentiation as seen in the developing brain. In addition, a small fraction of OLIG2^+^/GFP^+^ NPCs were able to generate cholinergic neurons identified by the expression of choline acetyltransferase (ChAT) in the organoids (Figures 1J, 1K). Taken together, these results show that the ventral forebrain organoid model in this study recapitulates the expression pattern of OLIG1 and 2 observed in human fetal brain tissue. Moreover, we provide direct evidence demonstrating that human OLIG2^+^/GFP^+^ NPCs give rise to different subclasses of GABAergic neurons and cholinergic neurons in 3D culture.

### Abnormal OLIG2 protein expression in DS hiPSC-derived ventral forebrain organoids

We next derived ventral forebrain organoids from the three pairs of control and hiPSCs generated in our previous study (Chen et al., 2014), including two DS hiPSC lines (DS1 and DS2) and isogenic disomic (Di)-DS3 and trisomic (Tri)-DS3 hiPSCs (Table S2 and Figures S2A, S2B). Control and DS pNPCs stably exhibited disomy and trisomy of HAS21, respectively (Figure 2A). At week 5, control and DS pNPCs similarly formed organoids with ventricular zone like regions containing Ki67^+^ and SOX2^+^ progenitors, and βIIIT^+^ immature neurons (Figure 2B). Similar expression of brain identity markers was also observed in control vs. DS organoids (Figures 2C, 2D). As expected, a significantly higher percentage of OLIG2^+^ cells was found in DS organoids, compared to control organoids (∼ 40% in control vs. 70% in DS, Figure 2E, 2F). Furthermore, we observed significantly higher expression of *OLIG2* mRNA (6 to14-fold), compared to control organoids (Figure 2G). This finding was verified by western blot analysis (Figures 2E, 2F). In addition, we also observed a higher expression of *OLIG1* transcripts in DS organoids (6 to 17-fold, Figure 2G). However, very few OLIG1^+^ cells were identified in either DS or control organoids by immunostaining and no significant differences were noted between the two groups (Figures 2C, 2D). Western blot analysis confirmed the similarly low expression of OLIG1 protein in both control and DS organoids (Figures 2H, 2I). These results further implicate the potential role of OLIG2 in neuronal differentiation in early stage NPCs.

**Figure 2.**
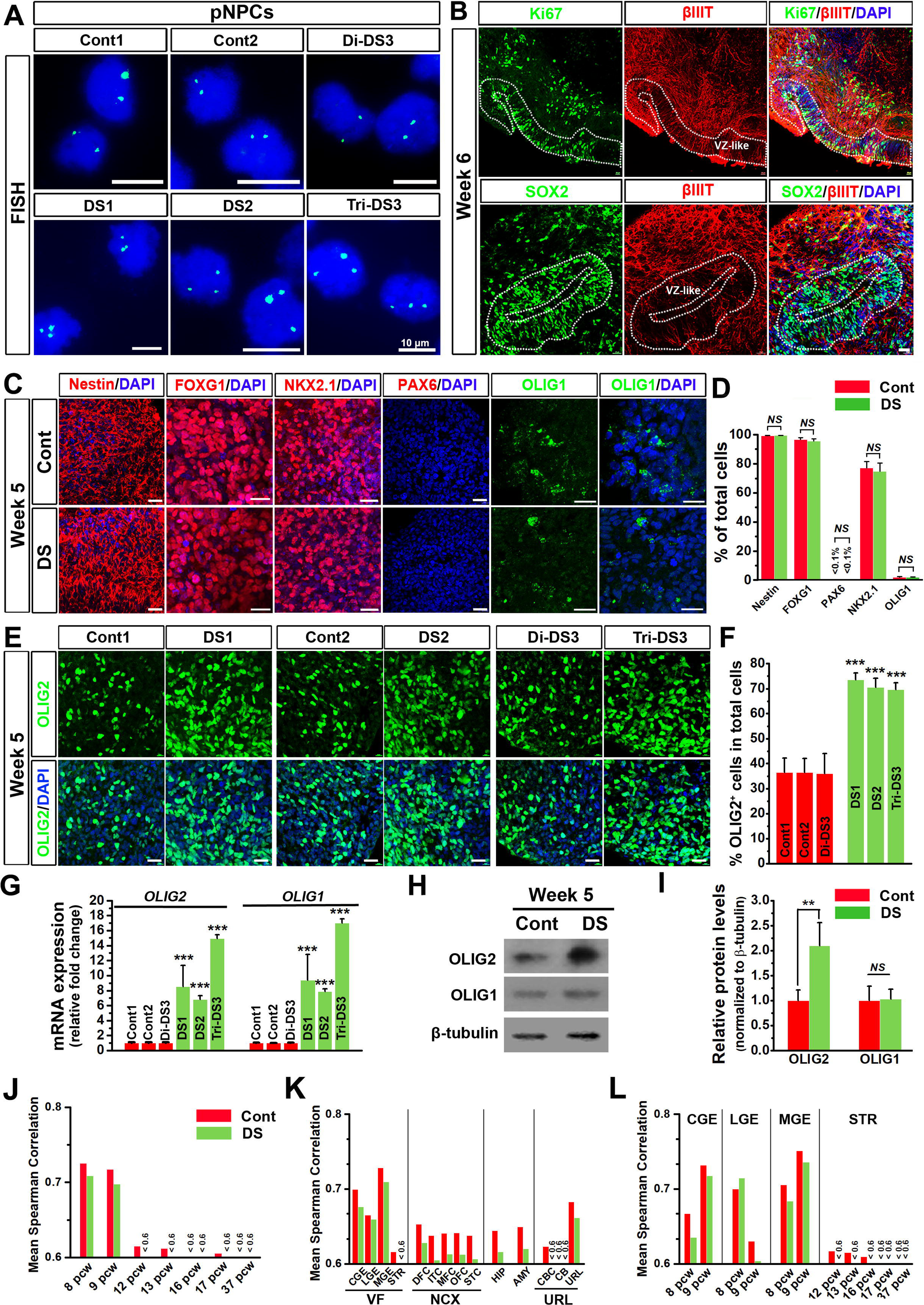
Abnormal expression of OLIG2 in DS ventral forebrain organoids. (A) Representatives of fluorescence in situ hybridization (FISH) analysis in the three pairs of control (Cont) and DS pNPCs. Scale bar represents 10 μm. (B) Representatives of Ki67, SOX2, and βIIIT immunostaining in 6-week-old control and DS organoids. VZ-like indicates ventricular zone-like area. Scale bars: 100 μm. (C and D) Representatives and quantification of Nestin-, NKX2.1-, FOXG1-, PAX6-, and OLIG1- expressing cells in 5-week-old control and DS organoids (n = 4). Scale bars: 20 μm. Student’s *t* test. *NS* represents no significance. (E and F) Representatives and quantification of OLIG2-expressing cells in 5-week-old control and DS organoids (n = 4). Scale bars: 20 μm. Student’s *t* test. \*\*\**P* < 0.001, comparison between DS and corresponding control organoids. (G) qPCR analysis of *OLIG2* and *OLIG1* mRNA expression in 5 to 6-week-old control and DS organoids (n = 4). Student’s *t* test, \*\*\**P* < 0.001, comparison between DS and corresponding control organoids. (H and I) Western blotting analysis and quantification of OLIG2 and OLIG1 expression in 5 to 6-week-old control and DS organoids (n = 3). Student’s *t* test. \*\**P* < 0.01. (J-L) Transcriptome correlation between 5-week-old organoids and post-mortem human brain samples from the BrainSpan project. The x axis shows the post-conception age in weeks or the brain region of the BrainSpan post-mortem brain samples. The y axis shows the mean spearman correlation. In (K) x axis shows BrainSpan agglomerated brain regions (i.e., more brain regions were merged into a single ‘‘larger’’ region): VF (i.e., VF, CGE, LGE, MGE, STR), ventral forebrain; NCX (i.e., DFC, ITC, MFC, OFC, STC), neocortex; HIP, hippocampus; AMY, amygdala; URL (i.e., CBC, CB, URL), upper rhombic lip. See also Figure S2, S5 and Table S2.

To assess the maturity and brain region specificity of the DS and control organoids, we obtained global transcriptome profiles of organoids by RNA-seq and compared them with BrainSpan, the largest dataset of postmortem human brain transcriptomes (Kang et al., 2011). Remarkably, the transcriptome analyses suggested that the hiPSC-derived 5-week-old organoids were most similar to the human brain at 8 to 9 weeks post-conception (pcw) (Figure 2J). Our preparation best reflected human ventral forebrain, including caudal ganglion eminence (CGE), lateral ganglion eminence (LGE), medial ganglion eminence (MGE), and striatum (STR), with smaller homology to the neocortex (NCX), hippocampus (HIP), amygdala (AMY), and upper rhombic lip (URL) (Figure 2K). We further comprehensively analyzed the maturity and ventral forebrain specificity of our organoids. Consistent results demonstrated that our preparation best reflected pcw 8 to 9 of MGE, CGE, and LGE, particularly the MGE. However, much less homology was found to STR that only appears after pcw 9 (Figure 2L).

### Overabundance of GABAergic neurons in DS organoids and early postnatal DS human brain tissue

To investigate GABAergic neuron generation, we cultured the control and DS organoids under the neuronal differentiation condition. Both DS and control organoids similarly increased in size over time in culture (Figures S2D, S2E). The NPCs in DS and control organoids efficiently differentiated to neurons; at week 8, we saw robust expression of MAP2 (Figure 3A). Similar percentages of NeuN^+^ neurons and S100β^+^ astrocytes were found (Figures 3B, 3C). Notably, GABA staining robustly labeled neuronal processes in addition to cell bodies in the organoids (Figure 3D, S2F). The GABA fluorescence intensity (FI) in the DS organoids was also significantly higher than that in control organoids (Figure 3E). More than half of the cells (∼60%) were GABA^+^ neurons in DS organoids, compared with only ∼35% in control organoids (Figures 3D, 3G).

**Figure 3.**
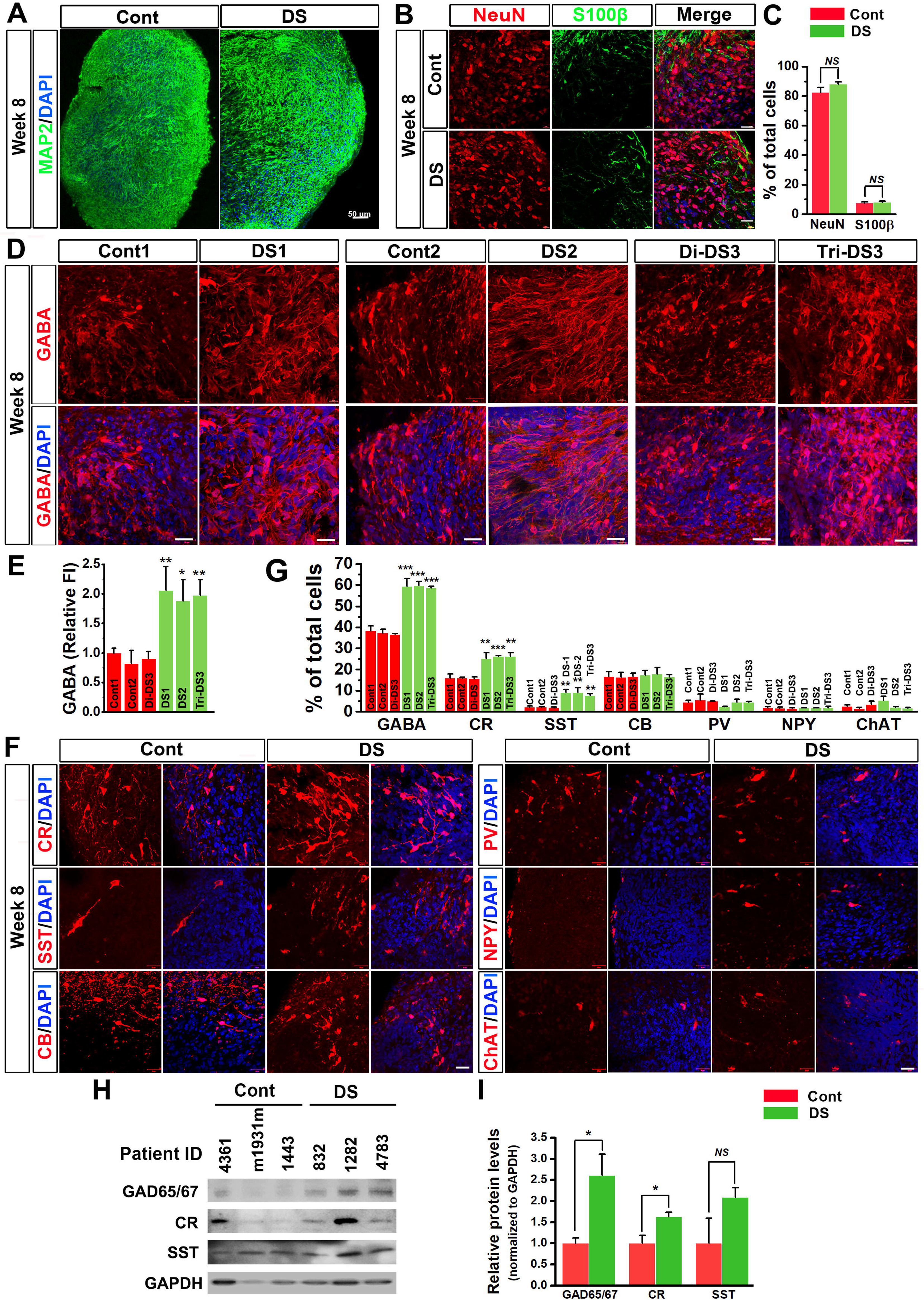
Overproduction of GABAergic neurons in DS organoids and DS human brain tissue. (A) Representatives of MAP2 expression in 8-week-old control and DS organoids. Scale bar: 50 μm. (B and C) Representatives and quantification of NeuN- and S100β-expressing cells in 8-week-old control and DS organoids (n = 4). Scale bar: 20 μm. Student’s *t* test. *NS* represents no significance. (D and E) Representatives of GABA-expressing neurons and quantification of fluorescence intensity (FI) of GABA staining in 8-week-old control and DS organoids (n = 4). Scale bar: 20 μm. Student’s *t* test. \**P* < 0.05, ** < 0.01, comparison between DS and corresponding control organoids. (F and G) Representatives and quantification of GABA-, CR-, CB-, SST-, PV-, NPY-, and ChAT- expressing neurons in 8-week-old control and DS organoids (n = 4). Scale bar: 20 μm. Student’s *t* test. \*\**P* < 0.01, \*\*\**P* < 0.001, comparison between DS and corresponding control organoids. (H and I) Western blotting analysis and quantification of GAD65/67, CR, and SST expression in postmortem cerebral cortex tissue from control and DS patients at ages less than one-year old. Student’s *t* test. \**P* < 0.05 and *NS* represents no significance. See also Figure S2 and Table S3.

In week 8 organoids, there were significantly more CR^+^ and SST^+^ GABAergic neurons in DS organoids than in control organoids (Figure 3F, 3G, S2F). There were no significant differences in the percentage of NPY^+^, CB^+^, and PV^+^ GABAergic neurons and ChAT^+^ cholinergic neurons (Figures 3F, 3G). To validate these findings, we examined the expression of CR, SST, and GAD65/67 in postmortem cerebral cortex tissue from DS and control subjects at ages of less than one year old (Table S3 and Figure S2G). We observed that protein levels of CR and GAD65/67 in DS patients, as compared to healthy controls, were significantly higher (Figures 3H, 3I). There was a trend of increased expression of SST in DS patients but with large variations, so no statistical significance was detected. This could be due to relatively fewer SST^+^ neurons in the cerebral cortex in neonatal human brain, compared to CR^+^ neurons (Paredes et al., 2016).

### DS neuronal chimeric mice show overproduction of GABAergic neurons

We further used a human neuronal chimeric mouse brain model (Chen et al., 2016) to examine the stability of the inhibitory neuronal fate of engrafted DS or control NPCs *in vivo* (Figure 4A). The donor-derived cells were tracked by staining human nuclei (hN) with specific anti-human nuclei antibody. Both engrafted DS and control cells were similarly distributed in mouse brains. At two months post-injection, the engrafted cells were mainly found near the injection sites (Figure 4B1), with some of the cells having migrated to the deep layers of cerebral cortex (Figure 4B2). Occasionally, a small cluster of cells was found in the superficial layers of cerebral cortex (Figure 4B2), which might have been deposited when the injection needle was retracted. The majority of hN^+^ donor-derived cells expressed NKX2.1 (Figures S3A, S3B). At six months post-transplantation, around 60% of donor-derived cells were LHX6^+^ in the cerebral cortex in both control and DS groups (Figures S3C, S3D), suggesting that the majority of the cells were post-mitotic interneuron precursors derived from the NKX2.1^+^ progenitors (Zhao et al., 2008). At six months post-injection, a large population of human cells dispersed widely in the cerebral cortex, as shown by hN staining in both sagittal and coronal brain sections (Figure 4C, S3E-S3K). Notably, there was a high density of donor-derived cells in the superficial layers and a low density in the deep layers (Figure 4C1), suggesting the migration of donor-derived cells from injection sites to the cerebral cortex. Doublecortin (DCX) is a marker of young migrating neurons (Gleeson et al., 1999). At 4 months post-injection, many hN^+^ cells expressed DCX and had bipolar processes (Figure S3L). We also tested injecting cells into deeper sites near the subventricular zone (SVZ) and found that the engrafted cells preferentially migrated along the rostral migratory stream to the olfactory bulb and did not lead to distribution and chimerization in the cerebral cortex (Figure S4). However, in contrast to our findings in the cerebral cortex, very few of these human cells in the olfactory bulb differentiated into CR^+^ cells (Figures S4C, S4F), suggesting that the vast majority of the neural progenitors generated using our current protocol were NKX2.1 lineage and gave rise to cortical or striatal interneurons, rather than olfactory bulb interneurons.

**Figure 4.**
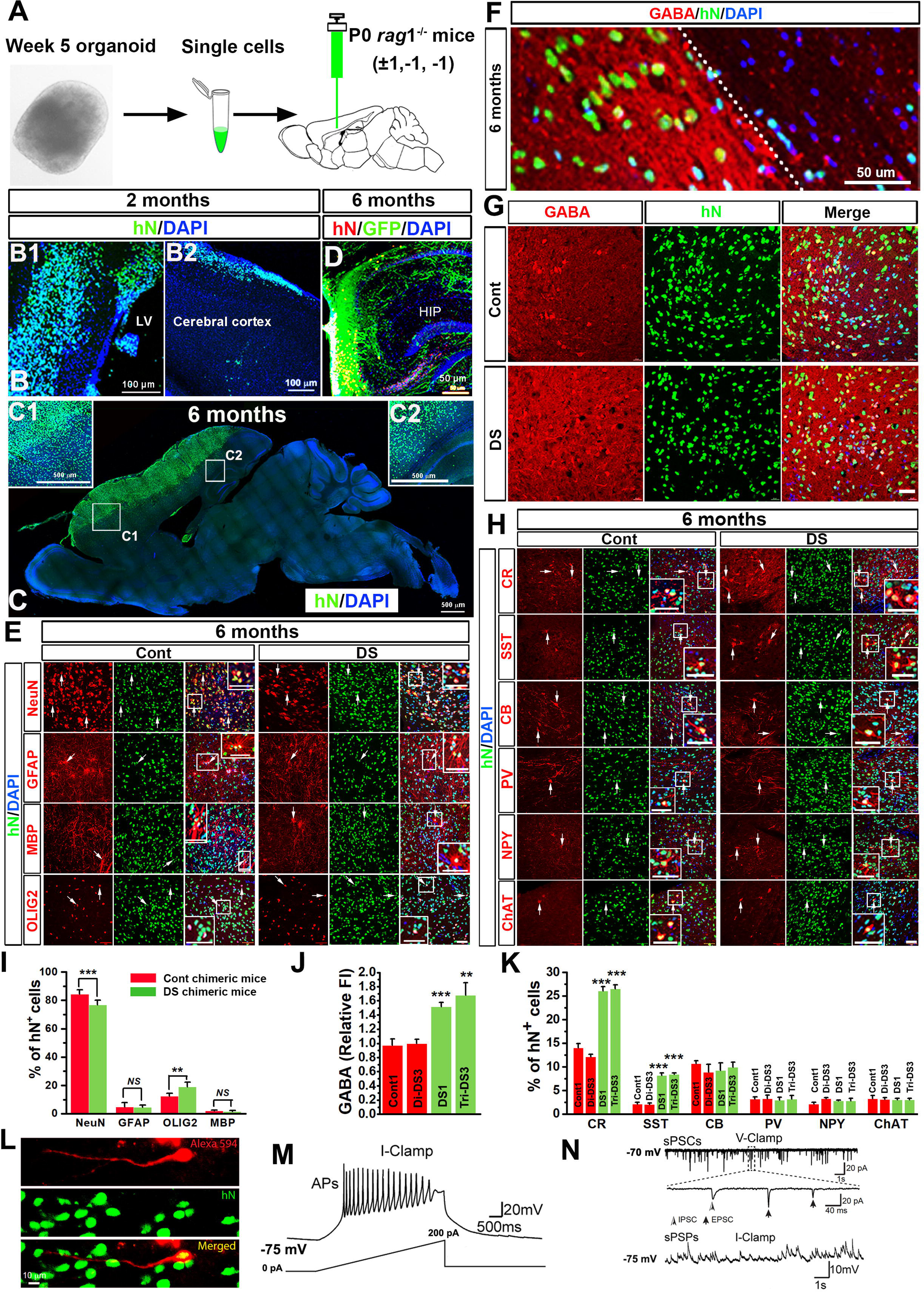
Human neuronal chimeric mice engrafted with DS ventral forebrain NPCs show overabundance of donor-derived GABAergic neurons and abnormal behavior. (A) A schematic diagram showing that organoids are dissociated into single cells and then engrafted into the brains of P0 *rag*1^−/−^ mice. (B) Representatives of hN^+^ engrafted human cells at two months post-transplantation. LV: lateral ventricles. Scale bars: 100 μm. (C) Representatives of hN^+^ cells at 6 months post-transplantation. The images are from confocal stitched tile scan. Scale bars: 100 μm. (D) A representative image of engrafted lenti-GFP-labeled cells. HIP: hippocampus. Scale bar: 50 μm. (E) Representatives of NeuN-, GFAP-, MBP-, and OLIG2-expressing cells differentiated from engrafted control and DS cells in the cerebral cortex at 6 months post-transplantation. Scale bars: 50 μm. (F) A representative image showing high FI of GABA staining in areas enriched of donor-derived hN^+^ cells. Scale bar: 50 μm. (G) Representatives of GABA staining in the cerebral cortex at 6 months after transplantation of control or DS cells. Scale bars: 50 μm. (H) Representatives of CR-, SST-, CB-, PV-, NPY-, and ChAT-expressing hN^+^ cells in the cerebral cortex at 6 months post-transplantation. Scale bars, 50 μm. (I) Quantification data showing the percentage of NeuN^+^, GFAP^+^, MBP^+^, and OLIG2^+^ cells among the total hN^+^ cells in the cerebral cortex at 6 months post-transplantation (n = 7 mice for each group). Student’s *t* test. \*\**P* < 0.01, \*\*\**P* < 0.001, and *NS* represents no significance. (J) Quantification data showing the FI of GABA staining in the cerebral cortex at 6 months post-transplantation (n = 7 mice for each group). Student’s *t* test. \*\**P* < 0.01, \*\*\**P* < 0.001, comparison between DS and corresponding control groups. (K) Quantification data showing percentage of CR^+^, SST^+^, CB^+^, PV^+^, NPY^+^, and ChAT^+^ cells among the total hN^+^ cells in the cerebral cortex at 6 months post-transplantation (n = 7 mice for each group). Student’s *t* test. \*\*\**P* < 0.001, comparison between DS and corresponding control groups. (L) Representatives of a human donor-derived neuron filled with Dextran Alexa Fluor 594 during patch-clamp recording and visualized by post hoc labeling with hN staining. Scale bars: 10 μm. (M and N) Representative current-clamp (I-clamp) and voltage-clamp (V-clamp) recording traces from the cell labeled in (L). APs: action potentials (APs); sPSCs: spontaneous postsynaptic currents; and sPSPs: spontaneous postsynaptic potentials. The enlarged area in the upper panel in (N) shows the presence of both excitatory and inhibitory postsynaptic currents, EPSCs and IPSCs, based on their respective kinetics. See also Figure S3, S4, S5, S6.

To specifically study the human GABAergic neurons developed in the context of cerebral cortex, we focused on analyzing the animals that had a wide distribution of donor-derived cells in the cerebral cortex at 6 months age. In some experiments, prior to transplantation, we labeled the cells with lentiviral-CMV-GFP and observed wide distribution of GFP^+^ processes in the hippocampus (Figure 4D). The control and DS cells were highly proliferative before transplantation, as demonstrated by high percentages of Ki67^+^ cells in both groups (Figures S5A, S5C). There were more OLIG2^+^/Ki67^+^ cells among total OLIG2^+^ cells in DS organoids than in control organoids (Figures S5A, S5C). After transplantation, the Ki67^+^/hN^+^ cells dramatically decreased from 2 months to 6 months (Figures S5B, S5D). Next, we analyzed differentiation of the engrafted cells in the cerebral cortex of 6-month-old chimeric mice. As shown in Figures 4E, 4I, a large proportion of the engrafted DS and control cells gave rise to NeuN^+^ neurons. The DS NPCs exhibited impaired neurogenesis, as indicated by a slightly reduced percentage of NeuN^+^/hN^+^ neurons among total hN^+^ cells in the DS cell transplantation group, as compared to the control cell transplantation group. A small fraction of the engrafted cells gave rise to GFAP^+^ astrocytes and MBP^+^ mature oligodendrocytes. No significant differences were noted in the percentages of astrocytes and mature oligodendrocytes in the two groups. Some of the donor-derived cells also expressed OLIG2, most of which at this stage were likely to be oligodendroglia progenitor cells, because no Nestin^+^ human NPCs were found in these 6-month-old chimeric mouse brains. We observed a higher percentage of OLIG2^+^/hN^+^ cells among total hN^+^ cells in the DS cell transplantation group than that in control cell transplantation group. We further examined the expression of OPC marker, PDGFRα. Consistently, there were a higher percentage of PDGFRα^+^/hN^+^ OPCs in DS group than in control group (Figures S5E, S5F). To examine GABAergic neuron differentiation from the engrafted cells, we double-stained the brain sections with GABA and hN. As shown in Figure 4F, S5G, and S5H, the areas that were strongly immunopositive for GABA staining were enriched for hN^+^ donor-derived cells, indicating the efficient differentiation of the engrafted cells to GABAergic neurons. These interneurons also expressed vesicular GABA transporter (VGAT) (Figure S5I). Engrafted DS cells overproduced GABAergic neurons, indicated by the much higher FI of GABA staining in DS cell transplantation group, as compared to control cell transplantation group (Figures 4G, 4J). Similar results were also obtained by examining the expression of GAD65/67 (Figures S6A, S6F). Furthermore, we found significantly higher percentages of CR^+^/hN^+^ and SST^+^/hN^+^ neurons among total hN^+^ cells in the DS group than in the control group (Figures 4H, 4K). No significant differences were noted in the percentage of CB^+^/hN^+^, PV^+^/hN^+^, and NPY^+^/hN^+^ GABAergic neurons, and ChAT^+^/hN^+^ cholinergic neurons (Figures 4H, 4K). Previous studies in mice show that a subpopulation of neocortical interneurons co-express CR and SST (Xu et al., 2006). We thus triple-stained control and DS chimeric mouse brains with CR, SST, and hN. At 6 months post-transplantation, none of the human interneurons in control chimeric mice co-expressed CR and SST, whereas in DS chimeric mice, CR^+^/SST^+^ human interneurons were occasionally seen (Figure S6B; less than 1% of total hN^+^ cells). These results suggest that the subclass of CR^+^/SST^+^ interneurons is increased in the DS chimeric mice, but they only represent a very small population. The overly produced interneurons from the engrafted DS cells are mainly CR^+^/SST^−^ and CR^−^/SST^+^ interneurons.

We further immunostained synaptic markers in the brain sections. As shown in Figure S6C, many of the hN^+^ cells in the cerebral cortex were surrounded by the presynaptic marker synapsin I. The PSD-95^+^ puncta, a postsynaptic marker, were found to distribute along the human-specific MAP2 (hMAP2)^+^ dendrites (Figure S6D). We then examined the expression of c-Fos, an activity-dependent immediate early gene that is expressed in neurons following depolarization and often used as a marker for mapping neuronal activity (Loebrich and Nedivi, 2009). As shown in Figures S6E, S6G, about 20% of the hN^+^ neurons at 6 months were positive for c-Fos, indicating that they were functionally active in the mouse brain. At 6 months post-transplantation, we performed electron microscopic analysis. We observed synaptic terminals formed between human neurons labeled by diaminobenzidine (DAB) staining against a human-specific cytoplasmic marker STEM121 and mouse neurons that were not labeled by the DAB staining (Figure S6H). In addition, we performed patch-clamp recording in brain slices and backfilled the recorded cells with fluorescent dye Dextran Alexa 594 through the recording pipettes. Among 22 recorded cells from 4 mice, 7 were identified as human neurons (Figure 4L). Out of the 7 hN^+^ donor-derived neurons, 6 neurons fired action potentials upon ramp current injections (Figure 4M) and all 7 neurons exhibited spontaneous postsynaptic currents (sPSCs) and potentials (sPSPs) (Figure 4N). These responses had sharp rises and slow decays, which are characteristics of synaptic events. In addition, both excitatory and inhibitory postsynaptic current responses were seen, based on their respective kinetics (Figure 4N, upper panel), suggesting that donor cell-derived GABAergic neurons might have interactions with host neuronal cells in the mouse brain.

### OLIG2 overexpression in DS biases gene expression attributed to GABAergic neurons

To gain mechanistic insight into how DS ventral brain NPCs were biased to GABAergic neuron generation, we performed RNA-sequencing using 5-week-old DS and control organoids. A dendrogram demonstrated that two control organoids and two DS organoids clustered closer to each other, respectively, indicating that they had similar biological properties within the same group, while the gene expression of DS organoids was distinct from that of control organoids (Figure 5A). As shown in Figures 5A, 5B, we began by comparing transcript expression between control1 and DS1 organoids. We identified a total of 2,920 DEGs, including 1,796 up-regulated genes and 1,124 down-regulated genes. We then examined the 2,920 DEGs in the organoids generated from isogenic Di-DS3 and Tri-DS3 hiPSCs. We identified a total of 610 overlapping DEGs, including 398 up-regulated genes and 212 down-regulated genes. Notably, there were 943 DEGs (506 up-regulated genes and 437 down-regulated genes) in the comparison between Di-DS3 and Tri-DS3 organoids that were not seen in the comparison between control1 and DS1 organoids. Next, we analyzed the distribution of the overlapping 610 DEGs on each human chromosome. As shown in Figure 5C, we found that the majority of the DEGs were non-HSA21 genes (577 of 610 DEGs). However, HSA21 had the highest percentage of DEGs among all the chromosomes. All of the DEGs on HSA21 were upregulated in DS samples and the overexpression of these genes on HSA21 resulted in both positive and negative regulation of the genes on other chromosomes. To further investigate how inhibiting the expression of *OLIG2* would change the global gene expression profile in DS during early embryonic brain development, we generated Tri-DS3 hiPSCs expressing OLIG2 shRNA (Tri-DS3 + OLIG2^shRNA^ hiPSCs). Then we derived organoids from the hiPSCs (Tri-DS3 + OLIG2^shRNA^ organoids). RNA-seq results revealed that many DEGs were reversed after OLIG2 knockdown, as compared to the organoids derived from the Tri-DS3 hiPSCs that received control shRNA with scrambled and non-targeting sequences (Tri-DS3 + Cont^shRNA^ organoids) and control1 organoids (Figure 5D). A Venn diagram showed that the expression of 255 of 398 upregulated overlapping DEGs and 166 of 212 downregulated overlapping DEGs was effectively reversed (Figure 5E). We also analyzed the reversed DEGs after OLIG2 knockdown compared between Di-DS3 and Tri-DS3 and their distribution on chromosomes (Figures S7A, S7B). Gene ontology (GO) analyses of the upregulated and downregulated genes showed that biological processes such as myelin sheath, transcription, and cell proliferation were significantly enriched (Figures 5F, 5G). Interestingly, GO and top 20 annotated pathways revealed that an enrichment of downregulated genes in DS organoids was particularly related to neurodevelopmental terms, indicating that the embryonic neuronal development as early as 8 to 9 pcw is altered in DS (Figure 5F). In addition, we found that the upregulated genes in DS organoids were enriched in extracellular matrix terms, indicating that the altered cells’ dynamic behavior in DS might affect cell adhesion, communication, and differentiation and fate specification (Figure 5G). Moreover, the GO analyses demonstrated that most of the changed pathways in both DS1 and Tri-DS3 groups were also significantly rescued, suggesting that inhibiting OLIG2 expression in DS effectively reversed the tendency for abnormal neurodevelopment (Figures 5F, 5G).

**Figure 5.**
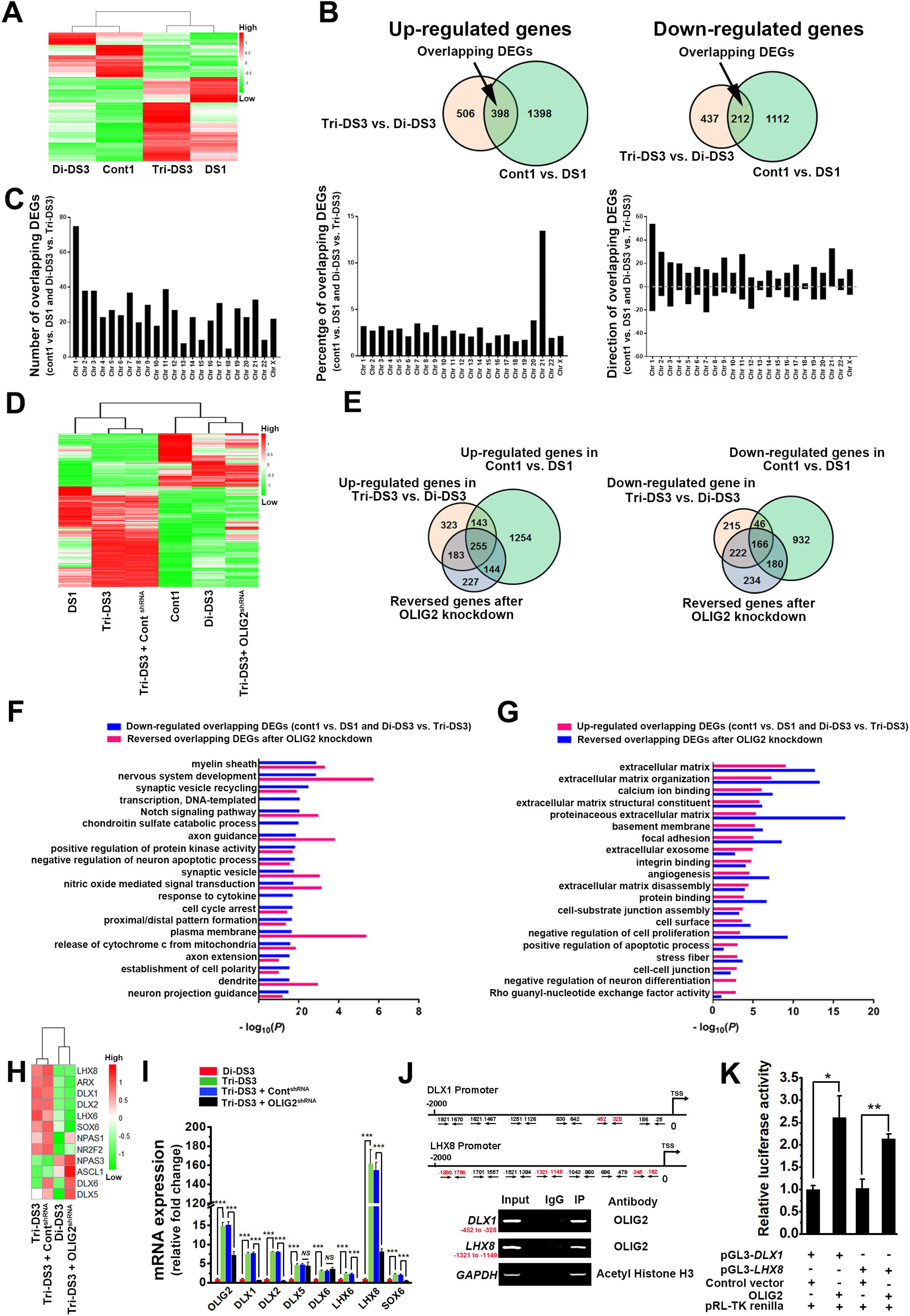
Abnormal gene expression in DS organoids. (A and B) A heatmap and Venn diagrams showing the differentially expressed genes (DEGs) in 5- week-old control and DS organoids. (C) The total number of DEGs, percentage of DEGs, and number of up-regulated or down-regulated DEGs on each chromosome in 5-week-old control and DS organoids. (D and E) A heatmap and Venn diagrams showing the rescued genes after OLIG2 knockdown. (F and G) GO analyses of the overlapping down-regulated (F) and up-regulated (G) DEGs, and the rescue effects after OLIG2 knockdown. (H) A heatmap showing the expression of genes critical for GABAergic neuron differentiation and specification in the 5-week-old organoids. (I) qPCR analysis of *OLIG2*, *DLX1*, *DLX2*, *DLX5*, *DLX6*, *LHX6*, *LHX8*, and *SOX6* mRNA expression in 5-week-old organoids (n = 4). One-way ANOVA test. \*\*\**P* < 0.001 and *NS* represents no significance. (J) ChIP assay showing that OLIG2 directly interacts with the promotor of *DLX1* and *LHX8*. The regions highlighted in red are the only ones that are immunoprecipitated by OLIG2 antibody. (K) Luciferase reporter assay showing that OLIG2 expression vector is capable of upregulating *DLX1* or *LHX8* luciferase activity, compared with control vector (n = 3). Student’s t test. * *P* < 0.01, ** *P* < 0.05. See also Figure S7 and Table S4-6.

RNA-seq results indicated that expression of many transcription factors with critical roles in GABAergic neuron development and specification was altered in DS organoids, as compared to control organoids. Among this group of transcription factors, *DLX1/2*, *LHX6/8*, *SOX6, and NPAS1/3,* which regulate interneuron fate (Azim et al., 2009; Flandin et al., 2011; Hanson et al., 2007; Stanco et al., 2014), were abnormally increased in the DS organoids. Other genes that regulate interneuron neurogenesis and migration, such as *ASCL1*(Casarosa et al., 1999; Castro et al., 2011), *ARX* (Colasante et al., 2008), and *NR2F2* (Kanatani et al., 2008) were also altered in DS organoids. Furthermore, we analyzed expression of these transcription factors after *OLIG2* knockdown. Strikingly, the majority of them were effectively reversed (Figure 5H). We further performed qPCR to verify RNA-seq results using brain organoids from the isogenic pair of hiPSCs. As shown Figure 5I, the expression of all the transcripts was increased in DS. Among all the genes we examined, only *DLX 5* and *6* showed inconsistent results between RNA-seq and qRT-PCR verification. This could be caused by the low expression levels of *DLX5* and *6,* because our qRT-PCR analysis indicated that *DLX5* and *6* were present at very low abundance in all the cell preparations. Moreover, the qPCR results demonstrated that inhibiting OLIG2 expression reversed the increased expression of the majority of the transcription factors (Figure 5I).

To further test our hypothesis that OLIG2 could regulate the expression of these transcriptional factors directly through interacting with the promoters of these genes, we performed a chromatin immunoprecipitation (ChIP) assay. We focused on examining the interactions between OLIG2 with *LHX8* or *DLX1* promoter sequences, because *LHX8* had the highest fold-change in the DS (Figure 5I) and *DLX1* was reported to be crucial for the production and longevity of CR^+^ and SST^+^ GABAergic neurons (Hanson et al., 2007). We designed 6 to 7 pairs of primers to span the 2000bp promoter sequences upstream of the *LHX8* and *DLX1* transcriptional start sites (TSS). As shown in Figure 5J, OLIG2 antibody immunoprecipitated the sequences highlighted in red in the *LHX8* promoter and *DLX1* promoter regions, demonstrating direct interactions of OLIG2 with promoters of *LHX8* and *DLX1*. We further performed dual luciferase reporter assay to examine the functionality of OLIG2 interactions with *DLX1* and *LHX8*. Based on our ChIP-PCR assay results, we cloned the promoter sequences of *DLX1* (−10 to −1250) and *LHX*8 (−1 to −2000) and inserted them into pGL3-Basic luciferase reporter vector. We then transfected these plasmids into control hiPSC-derived pNPCs. As shown in Figure 5K, when an OLIG2 expression vector was co-transfected into these cells, compared to co-transfection with a control vector, nearly 2.6 or 2.1-fold upregulation of luciferase activity was induced for *DLX1* or *LHX8*, respectively (Figure 5K).

### Inhibiting OLIG2 expression rescues overproduction of GABAergic neurons in DS organoids and chimeric mouse brains

To test our hypothesis that inhibiting the expression of OLIG2 would reverse the overproduction phenotypes under trisomy 21 conditions, we took the RNAi knockdown approach and used two DS hiPSC lines expressing OLIG2 shRNA or control shRNA (Figures S7C, S7D, S7E). We cultured the 5-week-old DS + OLIG2^shRNA^ and DS + Cont^shRNA^ ventral forebrain organoids under neuronal differentiation conditions and generated chimeric mouse brains by engrafting NPCs dissociated from these organoids. After 3 weeks (8-week-old organoids), we found that NeuN, S100β, PDGFRα, and NG2 expression were similar in disomic control, DS + OLIG2^shRNA^, and DS + Cont^shRNA^ organoids (Figures S7F, S7G). The percentages of GABA^+^ cells and FI of GABA staining were significantly decreased in DS + OLIG2^shRNA^ organoids, as compared to DS + Cont^shRNA^ organoids (Figures 6A, 6B). Moreover, the percentages of CR^+^ and SST^+^ GABAergic neurons were decreased after inhibition of OLIG2 expression (Figures 6C, 6D). The percentages of other subclasses of GABAergic neurons and cholinergic neurons were not significantly changed. In 6-month-old chimeric mouse brains, we found that the percentage of NeuN^+^/hN^+^ among total hN^+^ cells increased in the mouse brains which received DS + OLIG2^shRNA^ NPC transplant, as compared to the mouse brains which received DS + Cont^shRNA^ NPC transplant (Figures S7H, S7I). Consistent with the results in organoids, FI of GABA staining was decreased after inhibiting OLIG2 expression (Figures 6E, 6F). The overproduction of CR^+^ and SST^+^ GABAergic neurons was rescued and no significant differences in CB^+^, PV^+^, NPY^+^, and ChAT^+^ neurons populations were noted (Figures 6E and 6F).

**Figure 6.**
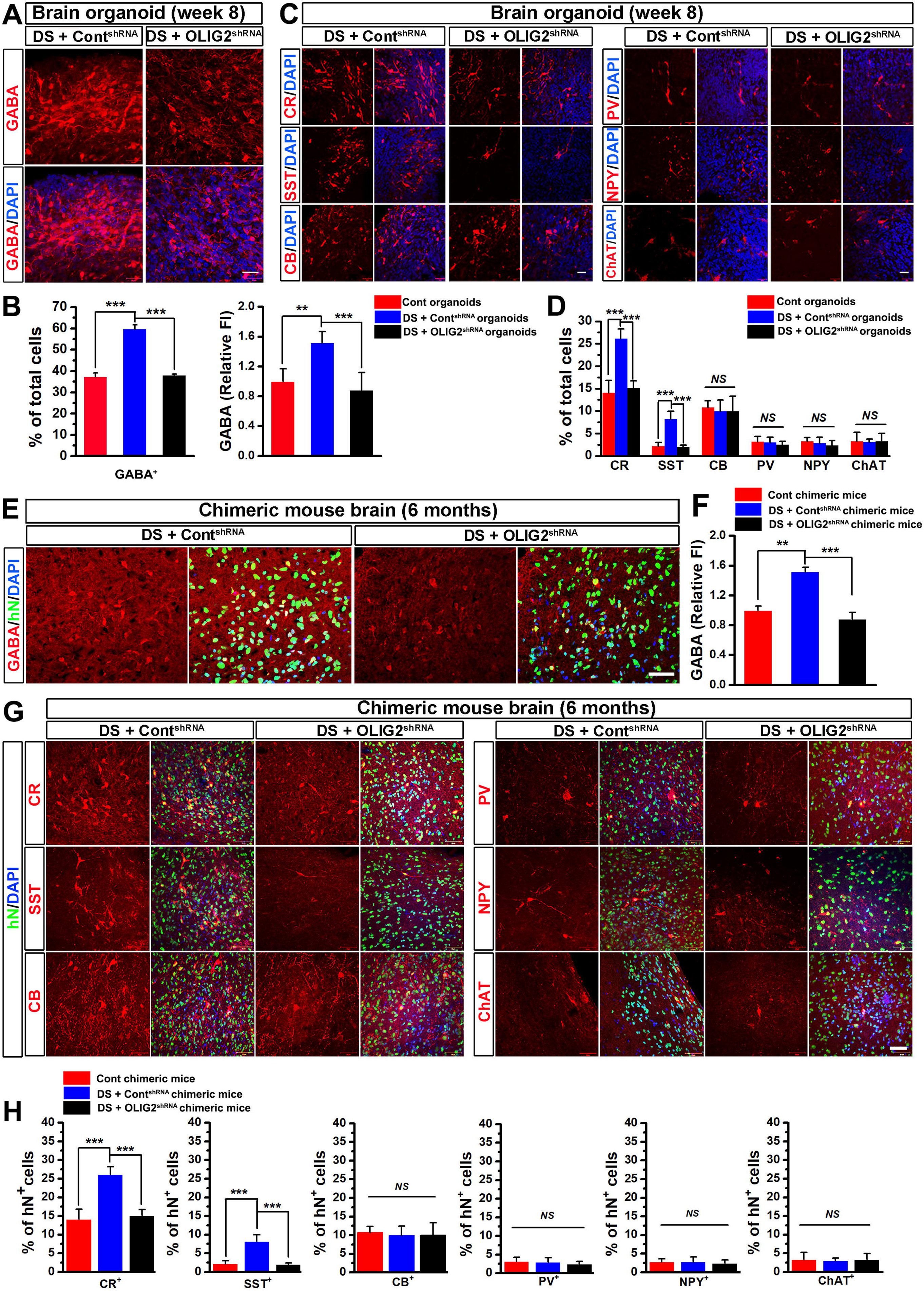
Inhibiting OLIG2 expression rescues overproduction of GABAergic neurons in DS organoids and chimeric mice. (A and B) Representatives and quantification of GABA^+^ cells and FI of GABA staining in 8-week-old disomic Cont organoids, DS + Cont^shRNA^ organoids, and DS + OLIG2^shRNA^ organoids (n = 4). Scale bars: 20 μm. One-way ANOVA test, \*\**P* < 0.01 and \*\*\**P* < 0.001. (C and D) Representatives and quantification of CR-, SST-, CB-, PV-, NPY-, and ChAT-expressing cells in 8-week-old Cont organoids, DS + Cont^shRNA^, and DS + OLIG2^shRNA^ organoids (n = 4). Scale bars: 20 μm. One-way ANOVA test, \*\*\**P* < 0.001, *NS* represents no significance (E and F) Representatives of GABA^+^ and hN^+^ cells and quantification of FI of GABA staining in the cerebral cortex at 6 months after transplantation of Cont, DS + Cont^shRNA^ or DS + OLIG2^shRNA^ cells (n = 6-8 mice for each group). Scale bars: 50 μm. One-way ANOVA test, \*\**P* < 0.01 and \*\*\**P* < 0.001. (G and H) Representatives and quantification of CR-, SST-, CB-, PV-, NPY-, and ChAT-expressing donor-derived hN^+^ cells in the cerebral cortex at 6 months after transplantation of Cont, DS + Cont^shRNA^ or DS + OLIG2^shRNA^ cells (n = 6-8 mice for each group). Scale bars: 50 μm. One-way ANOVA test, \*\**P* < 0.01 and \*\*\**P* < 0.001. See also Figure S7.

### DS chimeric mice exhibit impaired recognition memory that can be improved by inhibiting OLIG2 expression

Based on the findings that the donor cell-derived neurons might interact with host neurons in the mouse brain, we hypothesized that engrafted control and DS cells might differently impact the behavioral performance of the animals and that inhibiting OLIG2 expression in DS cells would reverse the different behavior phenotypes. We performed the open field test to evaluate basal global activity of the mice. Mice were placed into a chamber under dim ambient light conditions and activity was monitored for 5 minutes. We found no significant difference in the total traveling distance and time in the center area (Figure 7A; n = 10 - 15), suggesting no significant difference in locomotor activity and exploratory behavior among the five groups of mice. Next, we performed the novel object recognition test to assess learning and memory associated with the prefrontal cortex and hippocampus (Antunes and Biala, 2012; Wan et al., 1999). Similarly, no difference in the total traveling distance was observed (Figure 7B). We did not observe any significant behavioral changes in the preference ratio to the novel object between the PBS control group and control chimeric mouse group (Figure 7B). Interestingly, DS chimeric mice spent less time with the novel object, compared to control chimeric mice and PBS control mice (Figure 7B), suggesting that DS chimeric mice had impaired recognition memory. Moreover, this impairment was effectively rescued by inhibiting OLIG2 expression (Figure 7B). Next, we performed the elevated plus maze test to assess anxiety of the chimeric mice to further rule out the effects of anxiety on learning performance. We did not observe any significant difference in the total traveling distance, the number of entries, and time spent in the open arms (Figure 7C; n = 10-15), suggesting that anxiety was similar among the five groups. Moreover, we performed histological assessment after behavioral testing and found similar degrees of chimerization among the groups of animals that received cell transplant (Figure S7J).

**Figure 7.**
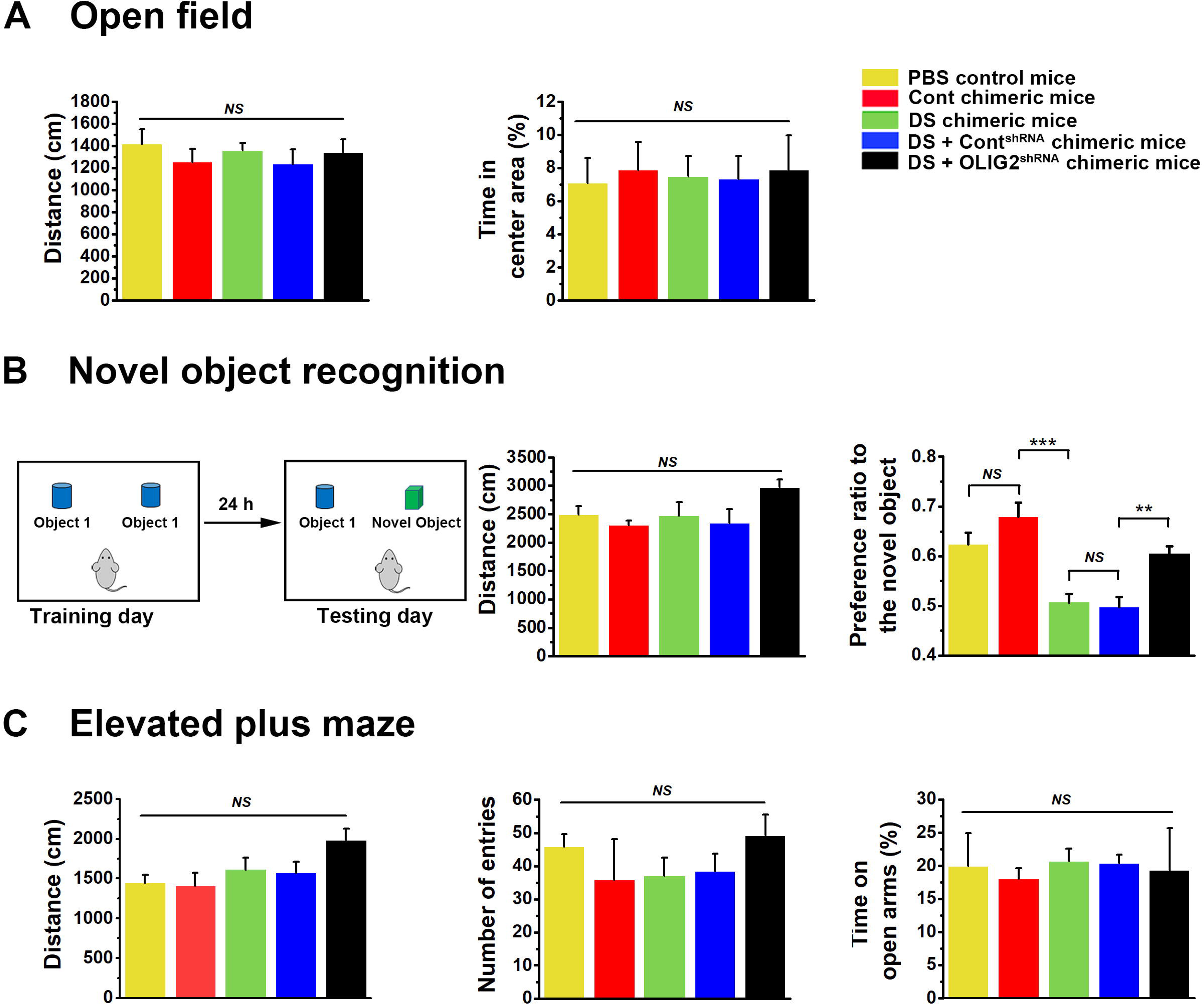
DS chimeric mice exhibit impaired cognitive functions. (A) Open field test showing the traveling distance and the percentage of time that PBS control mice, Cont chimeric mice, DS chimeric mice, DS + Cont^shRNA^ chimeric mice, and DS + OLIG2^shRNA^ chimeric mice spend in the center area of (n = 10-15 mice for each group). One-way ANOVA test. *NS* represents no significance. (B) *Left,* a schematic diagram showing novel object recognition test. *Right,* quantification of the traveling distance and the preference ratio to the novel object of the five groups of chimeric mice (n = 10-15 mice for each group). One-way ANOVA test. \*\*\**P* < 0.001, \*\**P* < 0.01. *NS* represents no significance. (C) Elevated plus-maze test showing the traveling distance, the number of entries, and the time that the chimeric mice spend on open arms (n = 10-15 mice for each group). One-way ANOVA test. *NS* represents no significance.

## DISCUSSION

In this study, we employ a complementary combination of human iPSC-derived cerebral organoid and chimeric mouse brain model systems to precisely model DS. The 3D cerebral organoid model possesses significant advantages for modeling human embryonic ventral forebrain development and neuronal differentiation. The ventral forebrain organoids closely recapitulate the expression pattern of OLIG1 and OLIG2 in the human embryonic brain under healthy and DS disease states. As opposed to 2D cultures, the OLIG2^+^ NPCs in the 3D cultures are capable of generating distinct subclasses of neurons. In addition, the comparison between our organoid samples and the BrainSpan dataset confirms that the gene expression patterns in the organoids exhibit similarities with human embryonic brain at 8 to 9 pcw (equivalent to GW 10 to 11) that precede fetal stages (Engle et al., 2004). Thus, these organoids provide unprecedented opportunities to investigate DS human brain development at embryonic stages. The development of the human chimeric mouse model permits the study of human interneuron development and disease pathogenesis *in vivo.* Recent studies in human brain tissue have demonstrated the widespread and extensive migration of immature neurons for several months in the neonatal brain (Paredes et al., 2016; Sanai et al., 2011). In particular, a study identified that these immature neurons are mainly young CR^+^ GABAergic neurons, which form an arching or “Arc” structure adjacent to the anterior body of lateral ventricle and within the neighboring subcortical white matter (Paredes et al., 2016). This “Arc” migratory stream takes the human GABAergic neurons to the frontal lobe of infants and young children (Paredes et al., 2016). In our study, at 2 months post-transplantation, most of the engrafted cells migrate away from injection sites and integrate into the brain regions overlying the lateral ventricles, which might mimic the “Arc” structure seen in the infant human brain. Very interestingly, at 6 months, the donor-derived cells, mostly CR^+^ GABAergic neurons, reside predominantly in the cerebral cortex. Notably, the majority of transplanted neural progenitors differentiated to neurons, with a small population giving rise to glial cells. We propose that the following two reasons may account for this observation. First, the engrafted neural progenitors have great neurogenic potential, because those organoid-derived neural progenitors are differentiated from rosette-type pNPCs which are highly neurogenic and stay at a primitive stage, as they are highly responsive to instructive neural patterning cues *in vitro* (Elkabetz et al., 2008; Li et al., 2011). Second, we transplant cells into the mouse brain at the earliest postnatal age, P0, as there is an age-related decline of the neurogenic niche in the brain through development (Katsimpardi et al., 2014; Villeda et al., 2011). The neurogenic niche in the neonatal mouse brain may promote neurogenesis from the transplanted cells, further facilitating neuronal differentiation (Chen et al., 2016).

In DS cells, the increased expression of *DLX1* may be largely responsible for the overproduction of CR^+^ interneurons, because previous studies in mice have shown that *Dlx1* is involved in production of CR^+^ interneurons (Hanson et al., 2007); and that increased expression of *LHX6/8* and *SOX6* may be largely responsible for the overproduction of SST^+^ interneurons, because *Lhx6/8* and *Sox6* regulate specification of SST^+^ interneurons (Azim et al., 2009; Flandin et al., 2011). Moreover, these transcription factors may also work together to fine-tune the production of different subclasses of interneurons, since previous studies have also shown that Dlx5/6 are directly downstream of Dlx1/2 (Anderson et al., 1997a; Anderson et al., 1997b; Zerucha et al., 2000) and Sox6 functions downstream of Lhx6 (Batista-Brito et al., 2009). Future gene network analysis of the RNA-seq data may help guide the investigation of the regulatory effects of OLIG2 on downstream interneuron transcriptional targets. In the chimeric mouse brain, we find that although a large number of human cells are seen, there is no significant behavioral changes between the PBS control and control chimeric mouse groups.

Our results show that only around 20% of the total donor-derived human cells are c-Fos^+^ and likely to be functionally active. Thus, the low percentage of active cells among the total control human interneurons may not be able to cause significant behavioral changes in the control chimeric mouse group, as compared to the PBS control group. Furthermore, DS chimeric mice exhibit impaired recognition memory in the novel object recognition test, compared to control chimeric mice and PBS vehicle control mice, despite the fact that similarly around 20% of total DS cells are c-Fos^+^ active interneurons. These data suggest that the impaired recognition memory in DS chimeric mice is very likely caused by the addition of these subclasses-specific human interneurons. This is further supported by our observations that inhibiting OLIG2 expression in DS cells corrects the abnormal production of the CR^+^ and SST^+^ interneurons and that DS+OLIG2^shRNA^ chimeric mice have improved recognition memory, compared to the DS+control^shRNA^ chimeric mice. Collectively, these results suggest that cognitive impairments in DS may be associated with the abnormal production of subclass-specific GABAergic neurons from OLIG2^+^ embryonic ventral forebrain NPCs.

We also discovered findings that are distinct from those identified previously in mice and in human samples. Studies using postmortem brain tissues from elderly DS patients (Kobayashi et al., 1990; Ross et al., 1984) and 2D cultures of DS iPSCs (Huo et al., 2018) showed reduced production of GABAergic neurons. The previous studies only used postmortem brain tissues from elderly patients with Alzheimer-like degeneration (Kobayashi et al., 1990; Ross et al., 1984), which may confound the evaluation of the cell counts of GABAergic neurons (Contestabile et al., 2017). A recent study (Huo et al., 2018) using DS hiPSCs reported discrepant findings that DS cells produced less CR^+^ neurons, although they similarly overproduced SST^+^ neurons. This discrepancy may result from the fact that 2D cultures were used in that study. In addition, the different transplantation strategies may also result in the discrepant *in vivo* findings. Our transplantation strategy was designed to mimic the GABAergic neuron development in an immature brain in the context of cerebral cortex, as opposed to the strategy of transplantation of young GABAergic neurons into a mature brain (8 to 10 weeks old mice) at sites very deep in the brain (medial septum), used in the recent study (Huo et al., 2018). In this study, although around 80% of engrafted human NPCs are NKX2.1^+^, we observe that in control chimeric mice, nearly all of the human donor-derived CR^+^ GABAergic neurons are CR^+^/SST^−^ and in DS chimeric mice, the vast majority of the overly produced CR^+^ neurons are also CR^+^/SST^−^. In contrast, in normal mouse brain, there are exceedingly few CR^+^/SST^−^ interneurons that are derived from Nkx2.1^+^ MGE progenitor cells, because the vast majority of these interneurons are CGE/non-Nkx2.1 lineage derived (Butt et al., 2005; Lopez-Bendito et al., 2004; Wonders and Anderson, 2006; Xu et al., 2004). This discrepancy may reflect a species difference, because a previous study shows that CR^+^/SST^+^ neurons are virtually absent in human brain and nearly all of the CR^+^ neurons are CR^+^/SST^−^ (Gonzalez-Albo et al., 2001). However, we also cannot exclude the possibility that OLIG2 overexpression in DS may result in conversion of MGE-like progenitors to CGE-like progenitors after the downregulation of Nkx2.1. Further studies are required to address this question.

## Acknowledgements

This work was in part supported by grants from the NIH (R21HD091512 and R01NS102382 to P.J. and an Institutional Development Award from the NIGMS under grant number P30GM110768 to P.J.), and Edna Ittner Pediatric Research Support Fund (to P.J.). Y.L. was supported by R01NS110707, Memorial Hermann Foundation-Staman Ogilvie Fund, the Bentsen Stroke Center, the TIRR Foundation through Mission Connect (014-115), and the Craig H. Neilsen Foundation (338617). R.P.H. was supported by R01ES026057, R01AT008458, and R21DA039686. We thank Dr. Alice Liu from Rutgers University and Dr. Guang Yang from Beckman Research Institute of City of Hope for suggestions and technical assistance with luciferase reporter assay. We also thank Mr. Andrew J. Boreland from Rutgers University for drawing the graphical abstract.

## Author Contributions

R.X. and P.J. designed experiments and interpreted data; R.X. carried out most of experiments with technical assistance from A.T.B., H.K., H.X., and W.K.; S.L., Y.L., and R.P.H. performed the gene expression analysis; J.L. and Z.P.P. performed electrophysiological recordings; Z.P.P. and R.P.H. provided critical suggestions to the overall research direction; P.J. directed the project and wrote the manuscript together with R.X. and input from all co-authors.

## Competing Financial Interests

The authors declare no competing financial interests.

## STAR METHODS

Detailed methods are provided in the online version of this paper and include the following:

- KEY RESOURCES TABLE
- CONTACT FOR REAGENT AND RESOURCE SHARING
- EXPERIMENTAL MODEL AND SUBJECT DETAILS

- Generation, culture, and quality control of hiPSC lines and their neural derivatives
- Animals
- METHOD DETAILS

- Cell transplantation
- Electrophysiology
- RNA isolation and qPCR
- ChIP
- RNA sequencing
- Immunostaining and cell counting
- Electron microscopy
- Western blotting
- Coomassie blue staining
- shRNA knockdown experiments
- FACS
- Cell fractionation
- Behavioral tests
- Open field test
- Elevated plus maze
- Novel object recognition
- Vector construction, transfection, and luciferase reporter assay
- CGH array.
- QUANTIFICATION AND STATISTICAL ANALYSIS
- DATA AND SOFTWARE AVAILABILITY SUPPLEMENTAL

## KEY RESOURCES TABLE

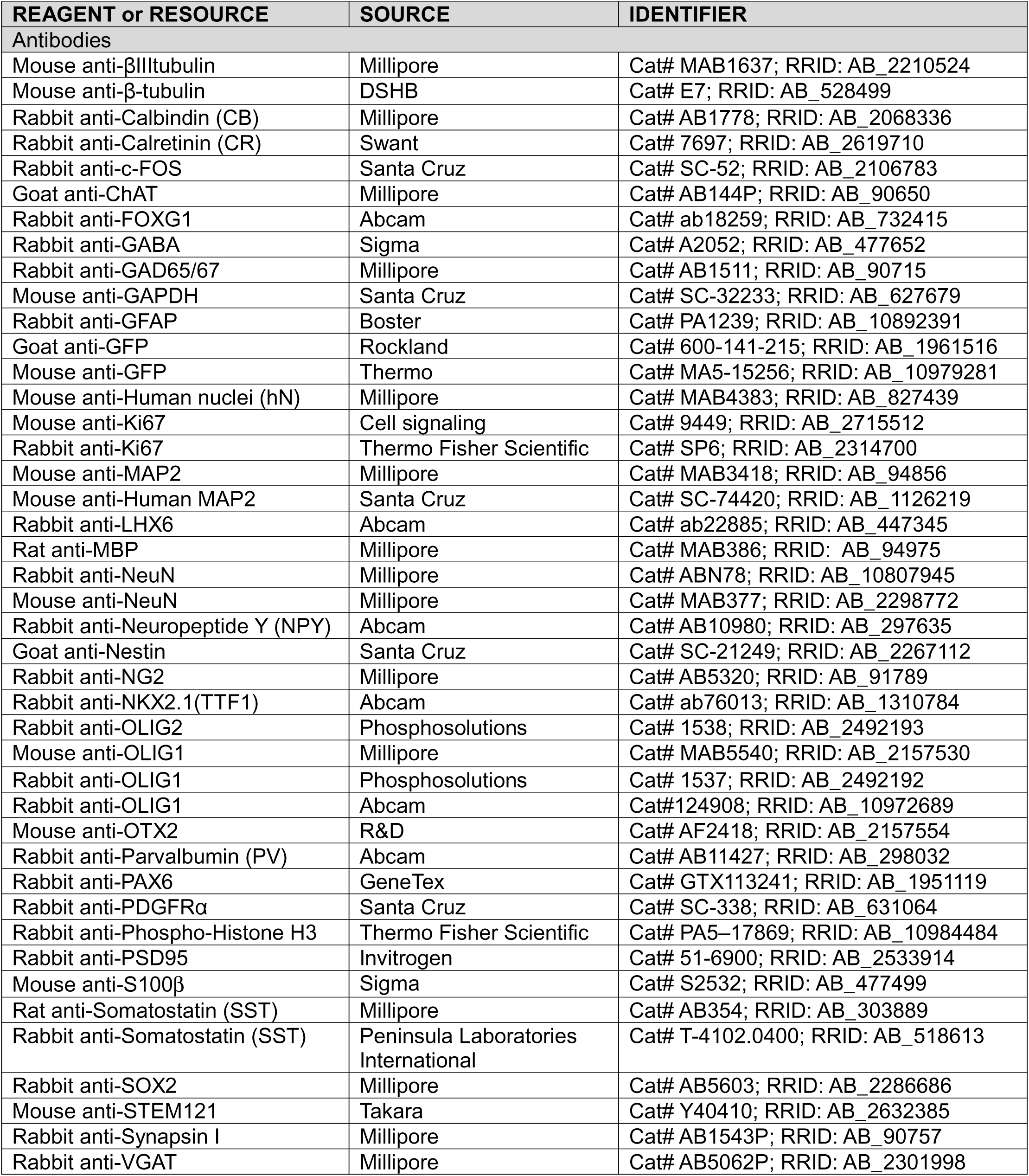
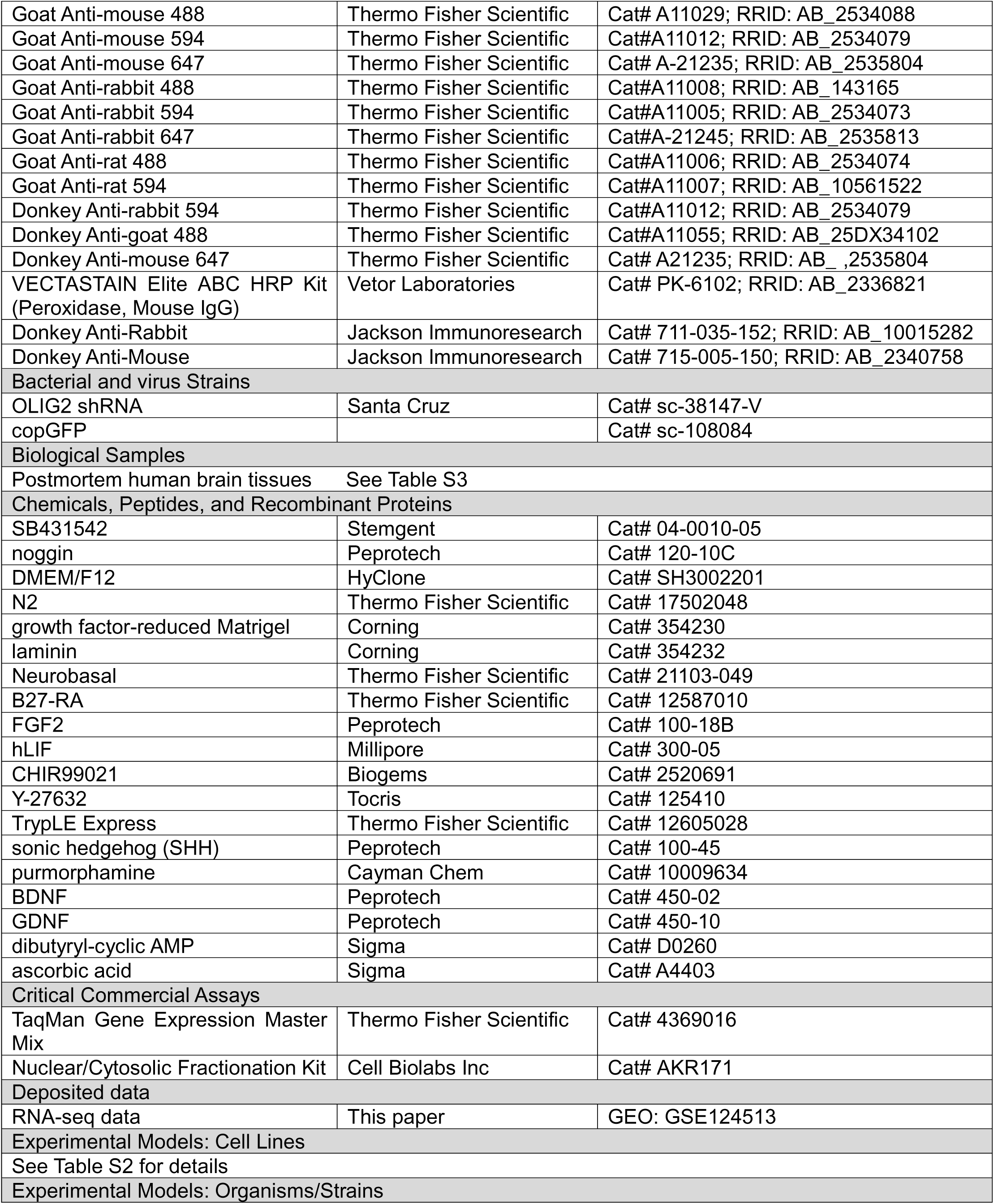
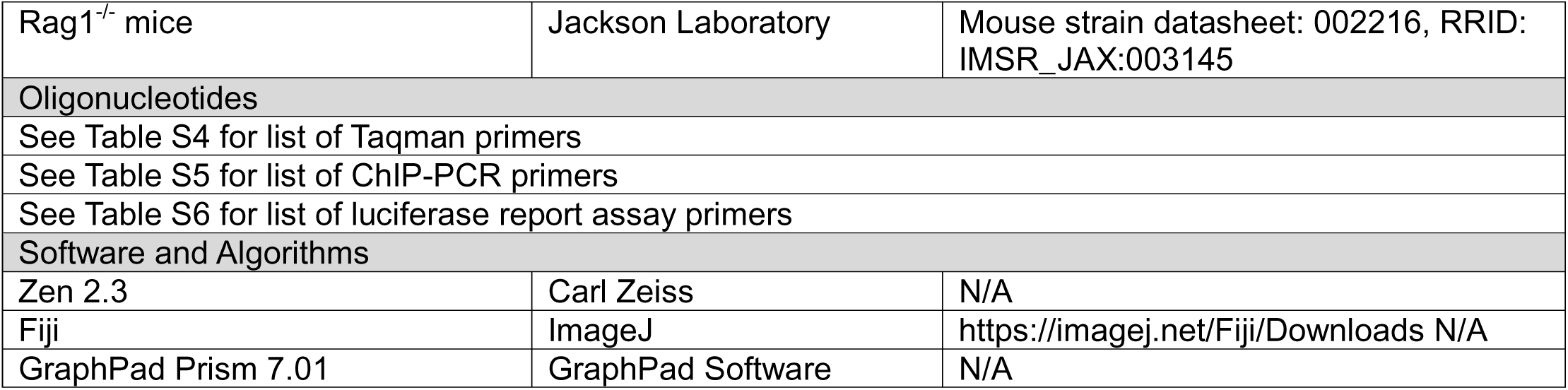

## CONTACT FOR REAGENT AND RESOURCE SHARING

Further information and requests for resources and reagents should be directed to and will be fulfilled by the Lead Contact, Peng Jiang.

## EXPERIMENTAL MODEL AND SUBJECT DETAILS

### Generation, culture, and quality control of hPSC lines and their neural derivatives

The control and DS hiPSC lines were generated from patient-derived fibroblasts using the “Yamanaka” reprogramming factors, as reported in our previous study (Chen et al., 2014). DS fibroblasts were obtained from Coriell Institute for Medical Research (Table S2). Totally, three pairs of control (Cont1, Cont2, and Di-DS3) and DS (DS1, DS2, and Tri-DS3) lines were used. The hiPSC lines derived from DS patients include two DS hiPSC lines (DS1, female; and DS2, male) and isogenic Di-DS3 and Tri-DS3 hiPSCs that were generated from a single female patient. As reported in our previous study (Chen et al., 2014), the same age-matched hiPSC lines (Cont1 and Cont2 hiPSCs) generated from healthy individuals were used as controls. Previous studies reported the generation of DS isogenic disomic and trisomic subclones either from the same parental iPSC lines during serial passage (Maclean et al., 2012) or from reprogramming mosaic DS fibroblasts (Weick et al., 2013). As reported in our previous study (Chen et al., 2014), we did not observe mosaicism in the fibroblasts from patient DS3 and the disomic subclones were identified during passaging similar to the study (Maclean et al., 2012). The isogenic Di- and Tri-DS3 iPSCs were then isolated and established by clonal expansion. Moreover, all the hiPSC lines derived from DS patients have been fully characterized by performing karyotyping, teratoma assay, DNA fingerprinting STR (short tandem repeat) analysis, gene expression profiling, and Pluritest (www.PluriTest.org), a robust open-access bioinformatic assay of pluripotency in human cells based on their gene expression profiles (Muller et al., 2011), as described in our previous study (Chen et al., 2014). The isogenic lines were verified by comparative genomic hybridization (CGH) array, which showed no significant insertions or deletions other than full HSA21 trisomy (Figures S2A, S2B). In addition, at the DNA level, all DS pNPCs consistently had ∼1.5-fold higher *OLIG1* and *OLIG2* gene copy numbers than the corresponding control pNPCs (Figure S2C). In the current study, fluorescence in situ hybridization (FISH) analysis with a human chromosome 21 (HSA21)-specific probe (Vysis LSI 21 probe; Abbott Molecular) was routinely performed to examine the copy number of HSA21 in the DS and control hiPSC-derived pNPCs, as previously described (Chen et al., 2014). In addition, *OLIG1* and O*LIG2* relative DNA copy number were examined routinely by using TaqMan Copy Number Assay (Applied Biosystems). Genomic DNA from DS and control pNPCs was collected by using DNA Extract All Reagents Kit (Applied Biosystems) and gene copy number of *OLIG1*, *OLIG2*, and RNase P were examined with primers Hs05549528, Hs02041572 or Copy Number Reference Assay human RNase P, respectively, using TaqMan Gene Expression Master Mix kit (Applied Biosystems). *OLIG1* and *OLIG2* relative DNA copy number were calculated by normalizing to RNase P DNA copy number, according to the manufacturer’s instructions. The OLIG2-GFP knockin hESC and hiPSC reporter lines were generated using a gene-targeting protocol as reported in our previous studies (Liu et al., 2011; Xue et al., 2009). The hPSCs were maintained under feeder-free condition and cultured on dishes coated with hESC-qualified Matrigel (Corning) in mTeSR1 media (STEMCELL Technologies). The hPSCs were passaged approximately once per week with ReLeSR media (STEMCELL Technologies). All the hPSC studies were approved by the committees on stem cell research at Rutgers University.

### Animals

Rag1^−/−^ mice (B6.129S7-*Rag1*^tm1Mom^/J, The Jackson Laboratory) were used for cell transplantation. All animal work was performed without gender bias under the Institutional Animal Care and Use Committee (IACUC) protocol approved by Rutgers University IACUC Committee.

## METHOD DETAILS

### Ventral forebrain organoid generation and culture

To generate ventral forebrain organoids, we used pNPCs as the starting population, because utilization of these neural fate-restricted progenitors has the advantage of efficient differentiation into desired brain regions (Monzel et al., 2017). Similar to previous studies (Mariani et al., 2012; Pasca et al., 2015; Xiang et al., 2017), we did not include Matrigel as supportive matrix in forming organoids, as we did not find significant improvement in the formation of ventral forebrain organoids by Matrigel embedding in our system. The pNPCs were derived from hPSCs using small molecule-based protocols (Chen et al., 2016; Li et al., 2011). Neural differentiation in the embryoid bodies (EB) was induced by dual inhibition of SMAD signaling (Chambers et al., 2009), using inhibitors SB431542 (5 μM, Stemgent) and noggin (50 ng/ml, Peprotech) in neural induction medium composed of DMEM/F12 (HyClone) and 1× N2 (Thermo Fisher Scientific) for 1 week. EBs were then grown on dishes coated with growth factor-reduced Matrigel (BD Biosciences) in the medium consisting of DMEM/F12, 1× N2, and laminin (1 μg/ml; Corning). pNPCs in the form of rosettes developed for another 7 days (week 2). Next, rosettes were manually isolated from surrounding cells and expanded for 7 days (week 3) in pNPC medium, composed of a 1:1 mixture of Neurobasal (Thermo Fisher Scientific) and DMEM/F12, supplemented with 1× N2, 1× B27-RA (Thermo Fisher Scientific), FGF2 (20 ng/ml, Peprotech), human leukemia inhibitory factor (hLIF, 10 ng/ml, Millipore), CHIR99021 (3 μM, Biogems), SB431542 (2 μM), and ROCK inhibitor Y-27632 (10 μM, Tocris).

Then the expanded pNPCs were dissociated into single cells using TrypLE Express (Thermo Fisher Scientific). 10,000 cells were cultured in suspension in the presence of ROCK inhibitor Y-27632 (10 μM) in ultra-low-attachment 96-well plates to form uniform organoids and then those organoids were transferred to ultra-low-attachment 6-well plates. To pattern these organoids to the fate of ventral forebrain, we treated them with sonic hedgehog (SHH) (50 ng/mL, Peprotech) and purmorphamine (1 µM, Cayman Chem), an agonist of sonic hedgehog signal pathway from week 3 to 5 (Figure 1A). The media were replenished every day. Starting from week 5, the cell culture plates were kept on orbital shaker with speed of 80 rpm/min. Maximum 8 organoids were cultured in each well of the ultra-low-attachment 6-well plate in neuronal differentiation medium containing a 1:1 mixture of Neurobasal and DMEM/F12, supplemented with 1× N2 (Thermo Fisher Scientific), 1× B27 (Thermo Fisher Scientific), BDNF (20 ng/ml, Peprotech), GDNF (20 ng/ml, Peprotech), dibutyryl-cyclic AMP (1mM, Sigma), and ascorbic acid (200 nM, Sigma) and media was replenished every other day. For 2D neuronal differentiation, 5,000 dissociated cells were plated onto poly-L-ornithine (0.002%, Sigma) and laminin (10 μg ml^−1^) pre-coated coverslips in the same neuronal differentiation medium. The neurons cultured in the neuronal differentiation medium for 4–6 weeks were used for experiments.

### Cell transplantation

For the transplantation experiments, two pairs of control and DS hiPSC lines, including control1 and DS1, and isogenic Di-DS3 and Tri-DS3 hiPSC lines, were used, because based on our results in organoids, all DS lines consistently showed similar trends of changes with statistical significance when compared with corresponding control lines (Figures 2G, 3E, 3G, 4J, and 4K). Moreover, inclusion of isogenic hiPSC lines can potentially limit the need for multiple iPSC lines to control for genetic variation (Chen et al., 2014; Maclean et al., 2012; Weick et al., 2013). The 5-week-old, patterned early ventral forebrain DS and control organoids were dissociated into single progenitor cells at a final concentration of 100,000 cells per μl in PBS. The cells were then injected into the brains of P0 *rag1* -/- immunodeficient mice (B6.129S7-Rag1tm1Mom/J on a C57BL/6 background, Jackson Laboratories). The precise transplantation sites were bilateral from the midline = ± 1.0 mm, posterior bregma = −1.0 mm, and dorsoventral depth = −1.0 or −1.8 mm (Figure 4A and Figure S4A). The mouse pups were anesthetized by placing them on ice for 5 minutes. Once cryo-anesthetized, the pups were placed on a digital stereotaxic device (David KOPF Instruments), equipped with a neonatal mouse adaptor (Stoelting). The pups were then injected with 1 μl of cells into each site by directly inserting Hamilton needles through the skull into the target sites. The pups were weaned at 3 weeks and were kept up to 6 months before they were tested for the engraftment of human cells. We did not observe any tumor formation of the transplanted cells in all the animals examined.

### Electrophysiology

Brain slice electrophysiology was carried out as described elsewhere (Liu et al., 2017). Mice were anesthetized, decapitated, and brains were removed and quickly immersed in cold (4°C) oxygenated cutting solution containing (in mM): 50 sucrose, 2.5 KCl, 0.625 CaCl_2_, 1.2 MgCl_2_, 1.25 NaH_2_PO_4_, 25 NaHCO_3_, and 2.5 glucose, pH to 7.3 with NaOH. Coronal cerebral cortex slices, 300 µm in thickness, were cut using a vibratome (VT 1200S; Leica). Brain slices were collected in artificial cerebrospinal fluid (ACSF) and bubbled with 5% CO_2_ and 95% O_2_. The ACSF contained (in mM): 125 NaCl, 2.5 KCl, 2.5 CaCl_2_, 1.2 MgCl_2_, 1.25 NaH_2_PO_4_, 26 NaHCO_3_, and 2.5 glucose (pH 7.3 with NaOH). After at least 1 hour of recovery, slices were transferred to a recording chamber and constantly perfused with bath solution (30°C) at a flow rate of 2 ml/min. Patch pipettes with a resistance of 8∼10 MΩ were made from borosilicate glass (World Precision Instruments) with a pipette puller (PC-10, Narishige) and filled with pipette solution containing (in mM): 126 K-Gluconate, 4 KCl, 10 HEPES, 4 Mg-ATP, 0.3 Na_2_-GTP, 10 phosphocreatine, (pH to 7.2 with KOH) for current and voltage clamp recordings. Dextran Alexa Fluor 594 was included in the intracellular recording solution for cell labeling. After whole-cell patch-clamp was achieved, spontaneous PSCs were recorded under voltage clamp at −70 mV. Input resistance and series resistance were monitored throughout the experiments and recordings were rejected if series resistance increased above 25 MΩ. All data were sampled at 5 kHz and analyzed offline using ClampFit 10.2 (Molecular Devices, USA) software. The identity of recorded cells was revealed by *post hoc* immunostaining with hN antibody and detected colocalization with the fluorescent dye.

### RNA isolation and qPCR

Total RNA was prepared from organoids with RNAeasy kit (Qiagen) (Chen et al., 2014). Complementary DNA was prepared with a Superscript III First-Strand kit (Invitrogen). The qPCR was performed with Taqman primers (Life Technologies) on an ABI 7500 Real-Time PCR System. All primers used are listed in Table S4. All experimental samples were analyzed and normalized with the expression level of housekeeping gene glyceraldehyde-3-phosphate dehydrogenase (GAPDH). Relative quantification was performed by applying the 2^−ΔΔCt^ method(Chen et al., 2014). For establishment of the ventral forebrain organoids model, qPCR analysis of *OLIG2* and *OLIG1* mRNA expression in the hPSC-derived 5-week-old ventral forebrain organoids (Figure 1E), the experiments are repeated for four times (n = 4) and for each experiment, 20 to 30 organoids derived from OLIG2- GFP hESCs and hiPSCs are used. For qPCR analysis of *OLIG2* and *OLIG1* mRNA expression in control and DS organoids (Figure 2G), three pairs of control (Cont1, Cont2, and Di-DS3) and DS (DS1, DS2, and Tri-DS3) iPSC-derived 5-week-old ventral forebrain were used. The experiments were repeated for four times (n = 4) and for each experiment, 20 to 30 organoids from each cell line were used. For validation the effect of *OLIG2* knockdown on transcription factor expression (Figure 5I), 5- week-old Di-DS3, Tri-DS3 organoids, and organoids derived from Tri-DS3 hiPSCs infected with lentivirus that carry control shRNA (Tri-DS3 + Cont^shRNA^) or OLIG2 shRNA (Tri-DS3 + OLIG2^shRNA^) were used, the experiments were repeated for four times (n = 4) and for each experiment, 20 to 30 organoids were used in each group.

### ChIP

Chromatin immunoprecipitation (ChIP) was performed using Magna-CHIP^TM^ A Chromatin Immunoprecipitation Kit (Millipore, USA) according to the manufacturer’s instructions. After being washed with cold PBS, the 5-week-old organoids were incubated in PBS with 1% formaldehyde at room temperature for ten minutes to cross-link chromatin proteins to genomic DNA. Glycine was used to quench unreacted formaldehyde. Following cold PBS washing, organoids were resuspended in lysis buffer with protease inhibitor Cocktail II and subjected to bioruptor sonication to shear DNA length to 200-1000 bp. 5 μg of anti-OLIG2 (Phosphosolutions 1538), anti-IgG (Kit included) or anti-Acetyl-Histone H3 antibody (positive control, included in the kit) and Protein A magnetic beads were added to the chromatin solution and incubated overnight at 4°C with rotation. After washing and elution, crosslinks of the protein/DNA complexes were reversed to free DNA by the addition of proteinase K and incubation at 62°C for 2 h followed by further incubation at 95°C for 10 min. Then, the DNA fragments were purified and used for PCR. The primers targeting human GAPDH was provided by the kit, working as the positive control. All primer sequences were listed in Table S5. The PCR products were analyzed by agarose gel electrophoresis.

### RNA sequencing

Organoids (5-week old) generated from DS hiPSC lines (DS1 and Tri-DS3) and control hiPSC lines (Cont1, isogenic Di-DS3), Tri-DS3+Cont^shRNA^, and Tri-DS3+OLIG2s^hRNA^ hiPSCs were used for RNA-sequencing. In order to reduce technical variations among different batches of cultures, a large number of organoids (total 40-60 organoids for each cell line) collected from three batches of organoid cultures were pooled together for RNA extraction and sample preparation.

Total RNA was prepared with RNAeasy kit (Qiagen) (Chen et al., 2014) and libraries were constructed using 600 ng of total RNA from each sample and the TruSeqV2 kit from Illumina (Illumina, San Diego, CA) following manufacturers suggested protocol. The libraries were subjected to 75 bp paired read sequencing using a NextSeq500 Illumina sequencer to generate approximately 30 to 36 million paired-reads per sample. Fastq files were generated using the Bc12Fastq software, version 1.8.4. The genome sequence was then indexed using the rsem-prepare-reference command. Each fastq file was trimmed using the fqtrim program, and then aligned to the human genome using the rsem-calculate-expression (version 1.2.31) command to calculate FPKM (fragments per kilobase of transcript per million mapped reads) values.

In order to analyze the transcripts, FPKM > 1 was set as cutoff to filter transcripts. |Fold change| > 1.5 was set as criteria to filter differential expressed genes (DEGs). We did two-step analyses to narrow down to the most prominent differentially DEGs that were identified in common between control1 vs. DS1 and Di-DS3 vs. Tri-DS3 organoids. Previous studies have shown that comparisons of RNA-seq data between isogenic pairs are often distinct from those between cell lines with different genetic backgrounds (Lin et al., 2015). Thus, choosing only transcripts overlapping between the two pairs of cultures provides an added level of confidence in their reproducibility.

Upregulated and downregulated DEGs were mapped to each chromosome and the number of DEGs in each chromosome was counted. Functional enrichment analysis was performed using DAVID online tool (Huang da et al., 2009a, b) and visualized using GraphPad. Heatmaps were generated using pheatmap package of R. Correlations with BrainSpan data were performed in R using only genes common to both experiments. The average of the Spearman correlation coefficient is presented for all the combinations of organoid sample replicates or BrainSpan samples matching the labeled condition.

RNA-seq experiments were performed by DNA Sequencing Core at UNMC, which receives partial support from the National Institute for General Medical Science INBRE - P20GM103427-14 and COBRE - 1P30GM110768-01 grants as well as The Fred & Pamela Buffett Cancer Center Support Grant - P30CA036727. We thank the Bioinformatics and Systems Biology Core at UNMC for providing RNA-seq data analysis services, which receives support from Nebraska Research Initiative and NIH (2P20GM103427 and 5P30CA036727).

### Immunostaining and cell counting

Mouse brains and organoids fixed with 4% paraformaldehyde were processed and cryo-sectioned for immunofluorescence staining (Liu et al., 2011; Pasca et al., 2015). The tissues were blocked with blocking solution (5% goat or donkey serum in PBS with Triton X-100) in room temperature (RT) for 1 hr. The Triton X-100 concentration was 0.1-0.2% for organoid and 0.8% for brain tissue. The primary antibodies were diluted in the same blocking solution and incubated with tissues in 4°C overnight. The sections were then washed with PBS and incubated with secondary antibodies for 1 hr in RT. After washing with PBS, the slides were mounted with the anti-fade Fluoromount-G medium containing 1, 40,6-diamidino-2-phenylindole dihydrochloride (DAPI) (Southern Biotechnology). The primary antibodies and secondary antibodies were listed in key resource table. Diaminobenzidine (DAB) immunostaining was performed with antibody a human-specific marker STEM121 as described in our previous study (Chen et al., 2016; Jiang et al., 2013b). Images were captured with a Zeiss 710 confocal microscope. Figure 4C, Figures S3E-S3J and Figure S4B scan images were obtained by confocal tile scan and automatically stitched using 10% overlap between tiles by Zen software (Zeiss). The analysis of fluorescence intensity was performed using ImageJ software (NIH Image). The relative fluorescence intensity was presented as normalized value to the control group. The cells were counted with ImageJ software. For organoids, at least three fields of each organoid were chosen randomly. For brain sections, at least five consecutive sections of each brain region were chosen. The number of positive cells from each section was counted after a Z projection. Engraftment efficiency and degree of chimerization were assessed by quantifying the percentage of hN^+^ cells among total DAPI^+^ cells in sagittal brain sections, as reported in the previous studies (Chen et al., 2015; Han et al., 2013). The cell counting was performed on every fifteenth sagittal brain section with a distance of 300 µm, covering brain regions from 0.3 to 2.4 mm lateral to the midline (seven to eight sections from each mouse brain were used). For the experiments using OLIG2-GFP hPSC reporter lines (Figure 1), 20 to 30 organoids derived from OLIG2-GFP hESCs and hiPSCs were used for each experiment and the experiments were repeated for four times (n = 4). For the comparison between control and DS organoids, 4 to 6 organoids from each cell line of the three pairs of control and DS were used for each experiment and the experiments were repeated for at least three times (n _>_ 3). For comparison between control and DS chimeric mouse brain, the data were generated from transplantation of cells derived from control (Cont1 and Di-DS3) and DS (DS1 and Tri-DS3) hiPSCs. For *OLIG2* shRNA knockdown organoids and chimeric mouse brain, we used control lines (control1, isogenic Di-DS3), DS lines (DS1 and Tri-DS3) and DS lines infected with lentivirus that carry control shRNA (DS1/Tri-DS3 + Cont^shRNA^) or OLIG2 shRNA (DS3/Tri-DS3 + OLIG2^shRNA^). The pool data of organoid experiments are repeated for four times (n = 4) and for each experiment, 4-6 organoids from each cell line were used. The pool data of chimeric mouse brain experiments were from transplantation of Cont, DS + Cont^shRNA^ or DS + OLIG2^shRNA^ ventral forebrain NPCs, 6-8 mice for each group.

### Electron microscopy

The 6-month-old mice were perfused with 0.1 M phosphate buffer (PB) followed by fixative solution (2% paraformaldehyde, 2.5% glutaraldehyde in 0.1 M PB pH 7.4). The brains were extracted and stored in the same fixative solution for 3 days at 4 °C. Then the brains were cut into 70 µm slices by vibratome. The sections were then subjected to Diaminobenzidine (DAB) immunostaining following the manual (Vector, USA) using antibody against a human-specific marker STEM121.The selected vibratome sections that contained DAB-labeled cells were post-fixed and processed for electron microscopy (EM) (Chen et al., 2016; Jiang et al., 2016; Jiang et al., 2013a). Brain sections were washed overnight in PBS, post-fixed for 2 hr at RT with 1.5% K_4_Fe(CN)_6_ and 1% OsO_4_ in 0.1 M PB, and then rinsed and stained with 3% uranyl acetate for 1 hr at 4°C. The sections were further transferred to 35%, 50%, 70%, 90% and 2 steps of 100% ethanol, and 2 steps of propylene oxide for dehydration. Next, the sections were embedded and sectioned on an ultramicrotome (Leica Ultracut UCT, Leica Microsystems). The collected sections were transferred onto grids and stained with uranyl acetate for 15-30 minutes and lead citrate for 3-15 minutes. EM images were captured using a high-resolution charge-coupled device (CCD) camera (FEI).

### Western blotting

Western blotting was performed as described previously (Xu et al., 2014). Lysates from organoids or human brain tissue were prepared using RIPA buffer and the protein content was determined by a Bio-Rad Protein Assay system. Proteins were separated on 4-12% SDS-PAGE gradient gel and transferred onto nitrocellulose membrane. Then the membrane was incubated with antibodies (Table S7). Appropriate secondary antibodies conjugated to HRP were used and the ECL reagents (Pierce ECL Plus Western Blotting Substrate, Thermoscientific) were used for immunodetection. For quantification of band intensity, blots from at least three independent experiments for each molecule of interest were used. Signals were measured using ImageJ software and represented as relative intensity versus control. β-tubulin was used as an internal control to normalize band intensity. For organoid western blotting analysis, the experiments are repeated for three times (n = 3) and for each experiment, 30 to 40 organoids respectively derived from three pairs of control and DS hiPSCs are used. The human brain tissues used for western blotting analyses are de-identified by encoding with digital numbers and obtained from University of Maryland Brain and Tissue Bank which is a Brain and Tissue Repository of the NIH NeuroBioBank. All the human brain tissues were derived from the cerebral cortex of patients at the ages from 186 to 339 days (Table S3). These human brain tissues were applicable for western blot analysis, since the denatured SDS-PAGE and Coomassie blue staining showed that the human brain sample preparations exhibited the absence of smear (little or no degradation) and sharp resolution of protein bands (Figure S2G). Additional information on the human brain tissue, including sex, race, and postmortem interval (PMI), was also provided.

### Coomassie blue staining

Similar to previous studies involving human tissue for western blotting (Blair et al., 2016), we performed denatured SDS-PAGE and Coomassie blue staining using the human samples, prior to examining the expression of proteins of interest. The sample preparations that exhibited the absence of smear (little or no degradation) and sharp resolution of protein bands were used for experiments. SDS-PAGE gradient gel was stained by staining solution (0.1% Coomassie Brilliant Blue R-250, 50% methanol and 10% glacial acetic acid) with shaking overnight and was de-stained using de-staining solution (40% methanol and 10% glacial acetic acid) for 1 hour.

### shRNA knockdown experiments

OLIG2 shRNA (sc-38147-V), non-targeting control shRNA (sc-108080), and copGFP (sc-108084) lentiviral particles were purchased from Santa Cruz Biotechnology. OLIG2 shRNA lentiviral particles carry three different shRNAs that specifically target *OLIG2* gene expression. DS1 and Tri-DS3 hiPSCs infected with lentivirus that carry control shRNA (DS1 or Tri-DS3 + Cont^shRNA^) or OLIG2 shRNA (DS1 or Tri-DS3 + OLIG2^shRNA^) were used in this study. Undifferentiated hiPSCs were infected with lentivirus and selected with puromycin treatment. Lentiviral particles in two different multiplicities of infections MOI=2 or 5) were mixed with 5 mg/ml polybrene in mTeSR and applied to hiPSCs overnight. Then, the infection medium was removed and replaced with fresh mTeSR. After three days culture, a puromycin selection was used. The optimal concentration of puromycin for the selection of transduced hiPSC colonies was established by puromycin titration on the specific hiPSC line used for the experiments. Puromycin was added at a concentration of 0.5 μg/ml for two weeks to select transduces hiPSCs. Stable hiPSCs were then used for neural differentiation with puromycin kept in all the media at 0.3 μg/ml as previously described (Mariani et al., 2015). Knockdown efficiency was confirmed by examining OLIG2 expression at both mRNA and protein levels after SHH and Pur treatment.

### FACS

5-week-old organoids derived from OLIG2-GFP hPSC, by treatment with DMSO or SHH and Pur, were dissociated to single cells by TrypLE (Gibico, 12605). About 1×10^7^ single cells suspended in PBS from each group were subjected to FACS each time. pNPCs without GFP signal were used as negative control. Sorting of OLIG2^+^/GFP^+^ cells and data acquisition were performed in a BD FACS Calibur (BD Bioscience). Acquired data were subsequently analyzed by calculating proportions/percentages and average intensities using the FlowJo software (Treestar Inc)(Jiang et al., 2013b). The purity of these cells was verified by immunostaining, showing that all of the sorted cells expressed GFP and OLIG2 (Figure S1D).

### Cell fractionation

The isolation of cell nuclear and cytosolic fractionation was performed by using Nuclear/Cytosolic Fractionation Kit (Cell Biolabs Inc, AKR171). The lysates of fractions were subjected to western blotting analyses. β-tubulin was used as the cytosolic marker and phospho-Histone H3 was used as the nuclear marker

### Behavioral tests

We examined five groups of mice: (1) vehicle control group that received injection of phosphate-buffered saline (PBS); (2) control chimeric mice that received transplantation of control1 or Di-DS1 cells; (3) DS chimeric mice that received transplantation of DS1 or Tri-DS3 cells; (4) DS + Cont^shRNA^ chimeric mice that received transplantation of DS1+Cont^shRNA^ or Tri-DS3+Cont^shRNA^ cells; and (5) DS + OLIG2^shRNA^ chimeric mice that received transplantation of DS1+OLIG2^shRNA^ or Tri-DS3+OLIG2^shRNA^ cells. We performed histological assessment after behavioral testing and assessed the engraftment efficiency and degree of chimerization by quantifying the percentage of hN^+^ cells among total DAPI^+^ cells in sagittal brain sections covering regions from 0.3 to 2.4 mm lateral to midline, as reported in the previous studies (Chen et al., 2015; Han et al., 2013). We found that the degree of chimerization was similar among the four groups of animals that received cell transplant (Figure S7J). Occasionally, animals that showed less than 3% chimerization were excluded from any analyses. All behavior tests were performed in a randomized order by an investigator blinded to treatment.

### Open field test

Mice were placed into a white Plexiglas chamber (60 long×60 wide×30 cm high) under dim ambient light conditions and activity was monitored for 5 min in a single trial. Four squares were defined as the center and the 12 squares along the walls as the periphery. The movements were monitored by an overhead video camera and analyzed by an automated tracking system (San Diego Instruments, CA). Results were averaged for total distance traveled and the number of entries into the center of the field (Belichenko et al., 2009; Ichiki et al., 1995).

### Elevated plus maze

The elevated plus maze was made of a black Plexiglas cross (arms 40 cm long × 6 cm wide) elevated 55 cm above the floor. Two opposite arms were enclosed by the transparent walls (40 cm long × 15 cm high) and the two other arms were open. A mouse was placed in the center of the apparatus facing an enclosed arm and allowed to explore the apparatus for 5 min. The total traveling distance and the time spent on the open arms was scored (Walf and Frye, 2007).

### Novel object recognition

As previously described (Belichenko et al., 2015; Bevins and Besheer, 2006), two sample objects in one environment were used to examine learning and memory with 24 hours delays. Before testing, mice were habituated in a black Plexiglas chamber (40 long×30 wide×30 cm high) for 10 minutes on 2 consecutive days under ambient light conditions. The activities were monitored by an overhead video camera and analyzed by an automated tracking system (San Diego Instruments, CA). First, two identical objects, termed as ‘familiar’, were placed in the chamber, and a mouse was placed at the mid-point of the wall opposite the sample objects. After 10 minutes to explore the objects, the mouse was returned to the home cage. After 24 hours, one of the ‘familiar’ objects used for the memory acquisition was replaced with a ‘new’ object similar to the ‘familiar’ one. The mouse was again placed in the chamber for 3 minutes to explore the objects; The total traveling distance and the time spent investigating the objects was assessed. The preference to the novel object was calculated as Time exploring new object / (Time exploring new object + Time exploring familiar object).

### Vector construction, transfection, and luciferase reporter assay

Promoter regions of human *DLX1* (from TSS upstream −10 to −1250 bp) or human *LHX*8 (from TSS upstream 0 to −2000 bp) were amplified by PCR from human genomic DNA extracted from control hiPSC-derived pNPCs with the primers shown in Table S6. Then promoter regions were inserted into the pGL3-Basic luciferase-reporter vector (Promega, USA). Control pNPCs were used for luciferase reporter assay. The pNPCs were seeded at 3X10^5^ per well in 24 well plates one day before transfection. The cells were co-transfected with 0.3 μg firefly luciferase-reporter construct (*DLX1*-pGL3, *LHX8*-pGL3 or pGL3-Basic), 0.02 μg pRL-TK Renilla luciferase-reporter plasmid (Promega, USA), and 0.6 μg pLenti-*Olig2-*IRES-GFP vector or control vector using Lipofectamine 3000 (Invitrogen, USA). The luciferase activity was examined using a dual-luciferase reporter assay system (Promega, USA) according to the manufacturer’s instructions, and the signal was normalized to the internal Renilla control for transfection efficiency.

### CGH array

CGH array was performed in the cytogenetic laboratory at Rutgers New Jersey Medical School (NJMS). The microarray CGH from NJMS is a customization of the Agilent Technologies 180,000 oligonucleotide probe array including SNPs to detect regions of homozygosity. These arrays provide genome-wide coverage of quantitative variants while as well as providing comprehensive assessment of known deletion/duplication syndromes and detailed coverage of all subtelomeric regions. Data were analyzed by BlueFuse Multi, v3.1 (BlueGnome, Ltd).

## QUANTIFICATION AND STATISTICAL ANALYSIS

All data represent mean ± s.e.m. The organoid experiments were repeated at least three times and each experiment was performed in triplicate using multiple stem cell lines. For immunohistochemical analysis, brain sections from at least 6-8 mice in each group were used. For behavior testing experiments,10-15 mice were used in each group. The data are shown as pooled results from DS and control lines or shown as results from individual pairs of DS and control lines. When only two independent groups were compared, significance was determined by two-tailed unpaired t-test with Welch’s correction. When three or more groups were compared, one-way ANOVA with Bonferroni post hoc test was used. A P value less than 0.05 was considered significant. The analyses were done in GraphPad Prism v.5.

## DATA AND SOFTWARE AVAILABILITY SUPPLEMENTAL

The accession number of the RNA sequencing data reported in this study is GEO: GSE124513.

## Notes

#### Summary of Updates

Figures and text have been revised

