## supplemental info for "OLIG2 Drives Abnormal Neurodevelopmental Phenotypes in Human iPSC-Based Organoid and Chimeric Mouse Models of Down Syndrome"

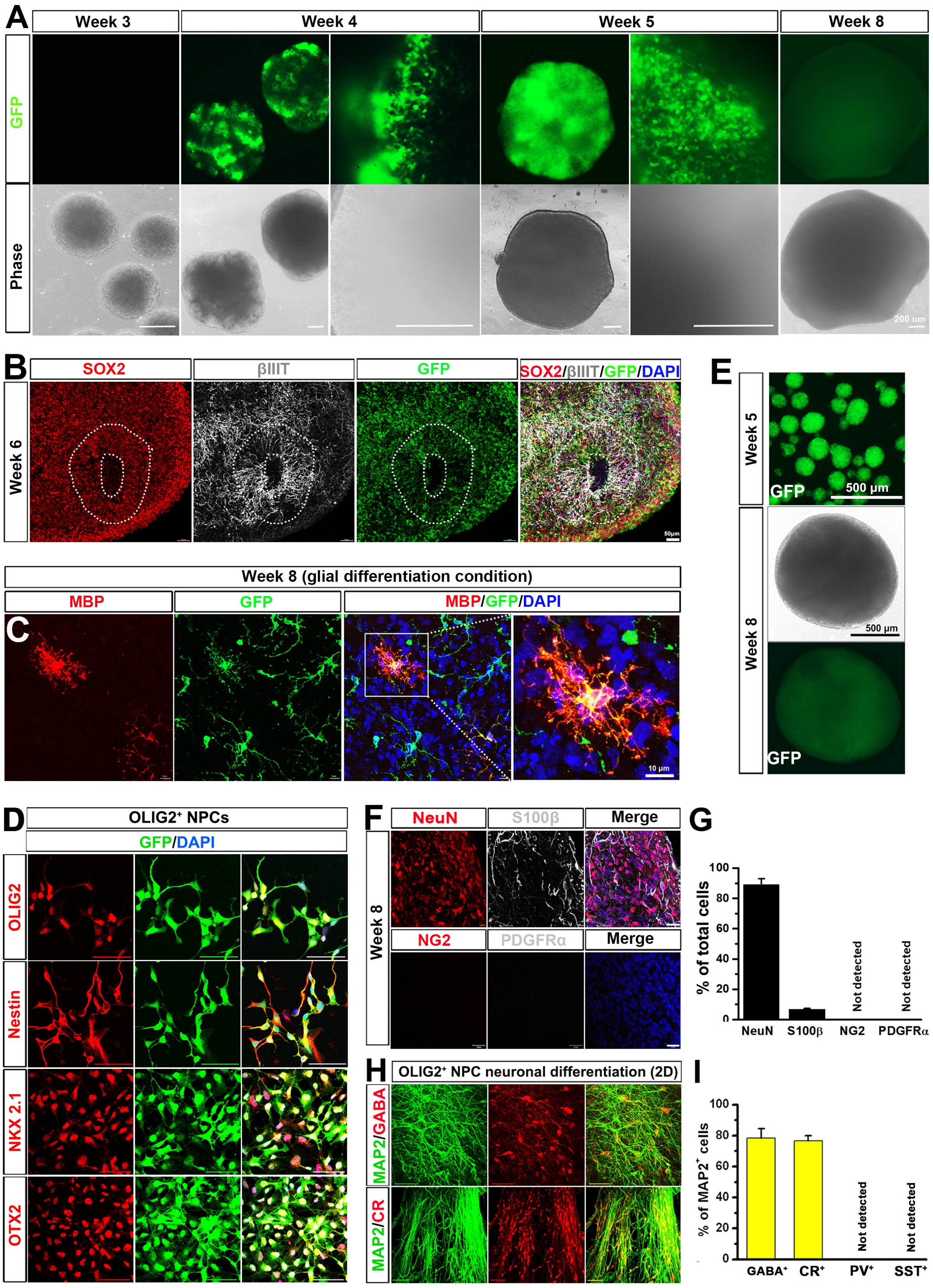
A
Week 3
Week 4
Week 5
Week 8
GFP
Phase
200 um
B
SOX2
βIIIT
GFP
SOX2/βIIIT/GFP/DAPI
Week 6
50μm
Week 8 (glial differentiation condition)
MBP
GFP
MBP/GFP/DAPI
C
10 μm
E
Week 5
GFP
500 μm
Week 8
500 μm
GFP
D
OLIG2+ NPCs
GFP/DAPI
OLIG2
Nestin
NKX 2.1
OTX2
F
NeuN
S100β
Merge
NG2
PDGFRα
Merge
Week 8
G
% of total cells
100
80
60
40
20
0
NeuN
S100β
NG2
PDGFRα
Not detected
Not detected
H
OLIG2+ NPC neuronal differentiation (2D)
MAP2/GABA
MAP2/CR
I
% of MAP2+ cells
100
80
60
40
20
0
GABA+
CR+
PV+
SST+
Not detected
Not detected

**Figure S1. Related to Figure 1. Ventral forebrain organoids derived from OLIG2-GFP hPSCs and $OLIG2^+$ ventral forebrain NPCs mainly give rise to $CR^+$ GABAergic neurons under 2D culture conditions.**

(A) Representative bright-field and fluorescence images of OLIG2-GFP hPSC-derived ventral forebrain organoids at different stages. Scale bar: 200 μm.

(B) Representatives of SOX2, βIIIT and GFP immunostaining, showing ventricular zone (VZ)-like areas in 6-week-old OLIG2-GFP hPSC-derived ventral forebrain organoids. Scale bars: 50 μm.

(C) Representatives of MBP-expressing mature oligodendrocytes in 8-week-old OLIG2-GFP hPSC-derived ventral forebrain organoids cultured under glial differentiation condition. Scale bars: 10 μm.

(D) Representatives of OLIG2-, Nestin-, NKX2.1-, and OTX2-expressing cells among the $GFP^+$ cells after FACS purification. Scale bars: 50 μm.

(E) Representative bright-field and fluorescence images showing $OLIG2^+$/$GFP^+$ cell-derived organoids at different time points. Scale bar: 500 μm.

(F) Representatives of NeuN-, S100β-, NG2-, and PDGFRα-expressing cells in 8-week-old $OLIG2^+$/$GFP^+$ cell-derived organoids cultured under neuronal differentiation condition. Scale bars: 20 μm.

(G) Quantification of pooled data from OLIG2-GFP hiPSCs and hESCs showing the percentage of $NeuN^+$, $S100\beta^+$, $NG2^+$, and $PDGFR\alpha^+$ in 8-week-old $GFP^+$ cell-derived organoids (n = 3 and for each experiment, 4 to 6 organoids from each line are used). Data are presented as mean ± s.e.m.

(H) Representatives of GABA-, CR-, and MAP2-expressing neurons differentiated from $OLIG2^+$ ventral forebrain NPCs under neuronal differentiation condition in 2D cultures. Scale bar: 50 μm.

(I) Quantification of pooled data from OLIG2-GFP hiPSCs and hESCs showing the percentage of $GABA^+$, $CR^+$, $PV^+$, and $SST^+$ cells among total $MAP2^+$ neurons (n = 4). Data are presented as mean ± s.e.m.

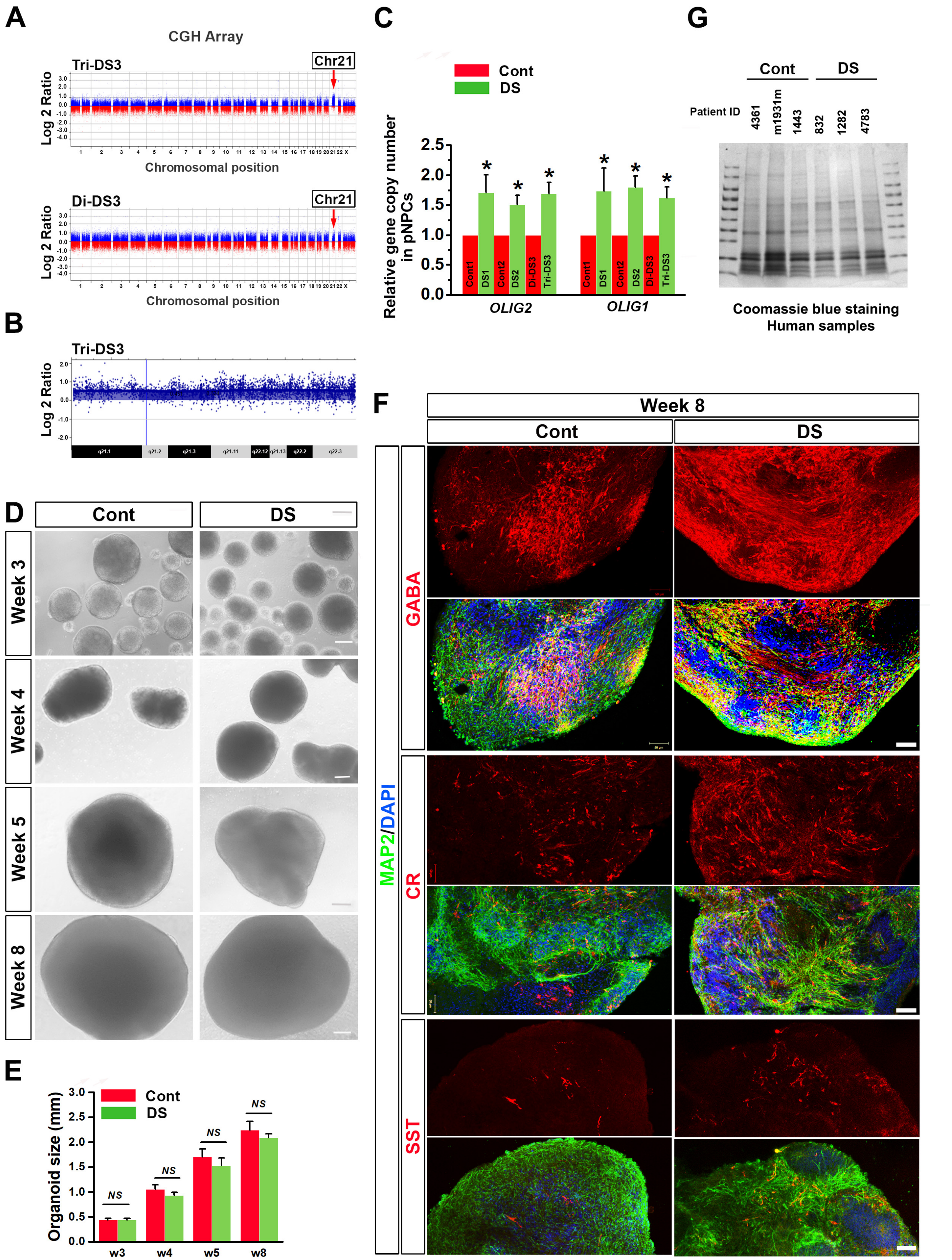
A
CGH Array
Tri-DS3
Chr21
Log 2 Ratio
Chromosomal position
Di-DS3
Chr21
Log 2 Ratio
Chromosomal position
B
Tri-DS3
Log 2 Ratio
C
Cont
DS
Relative gene copy number in pNPCs
Cont1
DS1
Cont2
DS2
Di-DS3
Tri-DS3
OLIG2
OLIG1
G
Cont
DS
Patient ID
4361
m1931m
1443
832
1282
4783
Coomassie blue staining
Human samples
D
Cont
DS
Week 3
Week 4
Week 5
Week 8
E
Organoid size (mm)
Cont
DS
NS
w3
w4
w5
w8
F
Week 8
Cont
DS
GABA
CR
SST
MAP2/DAPI

**Figure S2. Related to Figure 2 and 3. Overabundance of GABAergic neurons in DS organoids.**

(A) Comparative genomic hybridization (CGH) array of isogenic pair of Di-DS3 and Tri-DS3 demonstrates a gain of one chromosome 21 (red arrow) in Tri-DS3. No significant deletions or insertions were detected between the isogenic pair.

(B) Close-up of Chr21 CGH of Tri-DS3 shows full chromosome 21 trisomy with no deletions or duplications.

(C) qPCR results from genomic DNA showing the relative *OLIG1* and *OLIG2* DNA copy numbers in control (Cont1, Cont2, and Di-DS3) and DS (DS1, DS2, and Tri-DS3) pNPCs. $^*P < 0.05$, comparison between Cont1 vs. DS1, Cont-2 vs. DS2, or Di-DS3 vs. Tri-DS3 pNPCs.

(D) Representative bright-field images of control (Cont) and DS hiPSC-derived ventral forebrain organoids at different stages. Scale bars: 200 μm.

(E) Quantification of size of control (Cont) and DS hiPSC-derived ventral forebrain organoids at different stages. Scale bars: 200 μm. Student's *t* test. *NS* represents no significance. Data are presented as mean ± s.e.m.

(F) Representative low magnification images of organoids showing GABA-, CR- and SST-expressing neurons in 8-week-old control and DS organoids. Scale bar: 50 μm.

(G) Coomassie blue staining of the human samples after running denatured SDS-PAGE showing high sharpness and resolution of protein bands and the absence of smear (little or no degradation) of protein bands.

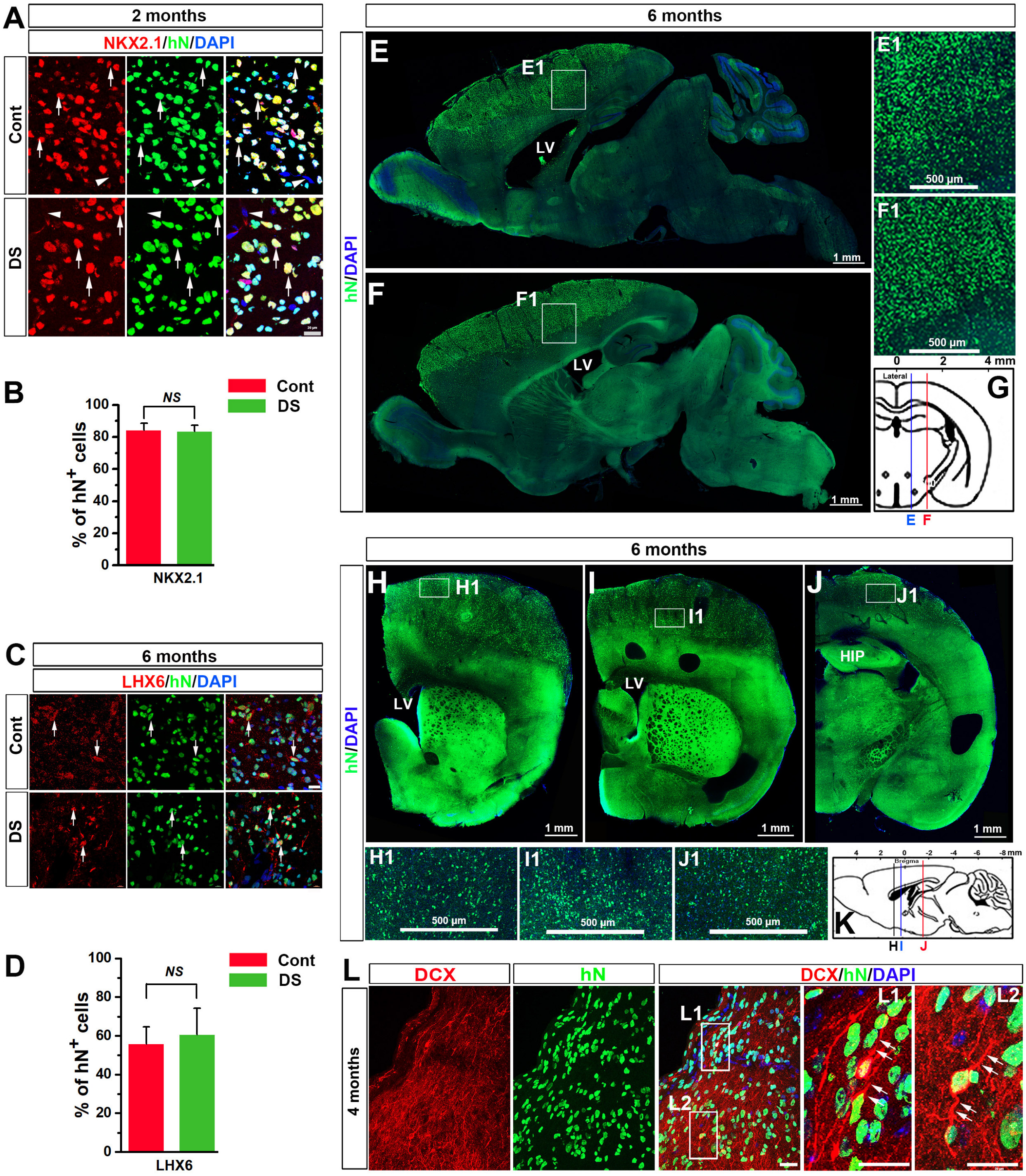
A
2 months
NKX2.1/hN/DAPI
Cont
DS
B
NS
Cont
DS
% of hN+ cells
100
80
60
40
20
0
NKX2.1
C
6 months
LHX6/hN/DAPI
Cont
DS
D
100
80
60
40
20
0
NS
Cont
DS
% of hN+ cells
LHX6
6 months
hN/DAPI
E
E1
LV
1 mm
F
F1
LV
1 mm
E1
500 µm
F1
500 µm
0
2
4 mm
Lateral
G
E
F
6 months
hN/DAPI
H
H1
LV
1 mm
I
I1
LV
1 mm
J
J1
HIP
1 mm
H1
500 µm
I1
500 µm
J1
500 µm
4
2
0
-2
-4
-6
-8 mm
Bregma
K
H
I
J
L
DCX
hN
DCX/hN/DAPI
4 months
L1
L2
L1
L2

**Figure S3. Related to Figure 4. Distribution and characterization of donor-derived cells after transplantation to the sites overlying lateral ventricles.**
(A) Representatives of NKX2.1-expressing donor-derived $hN^+$ cells in the cerebral cortex at 2 months post-transplantation of control or DS NPCs. Arrows indicate the donor-derived $hN^+$ cells expressing NKX2.1 and arrowhead indicates the NKX2.1-negative endogenous mouse cell. Scale bars, 20 μm.
(B) Quantification of pooled data from control (Cont1 and Di-DS3) and DS (DS1 and Tri-DS3) hiPSCs showing the percentage of $NKX2.1^+$ cells among the total $hN^+$ cells in the cerebral cortex at 2 months post-transplantation (n = 7 mice for each group). Student's *t* test. *NS* represents no significance. Data are presented as mean ± s.e.m.
(C) Representatives of LHX6-expressing donor-derived $hN^+$ cells in the cerebral cortex at 6 months post-transplantation of control or DS NPCs. Arrows indicate the donor-derived $hN^+$ cells expressing LHX6. Scale bars, 20 μm.
(D) Quantification of pooled data from control (Cont1 and Di-DS3) and DS (DS1 and Tri-DS3) groups showing the percentage of $LHX6^+$ cells among the total $hN^+$ cells in the cerebral cortex at 6 months post-transplantation (n = 7 mice for each group). Student's *t* test. *NS* represents no significance. Data are presented as mean ± s.e.m.
(E and F) Representative images from sagittal brain sections showing the wide distribution of $hN^+$ donor-derived cells in the cerebral cortex at 6 months post-transplantation. Areas in E1 and F1 are enlarged. The images were from confocal stitched tile scan. Scale bars: 1mm or 500 μm as indicated.
(G) A schematic diagram showing the relative positions of the sagittal sections shown in (E) and (F).
(H-J) Representative images from coronal brain sections showing the wide distribution of $hN^+$ donor-derived cells in the cerebral cortex at 6 months post-transplantation. The sections were collected after whole-cell patch-clamp recording and visualized by post hoc labeling with hN staining. Areas in H1, I1 and J1 are enlarged. The images were from confocal stitched tile scan. Scale bars: 1mm or 500 μm as indicated.
(K) A schematic diagram showing that relative positions of coronal sections shown in (H), (I), and (J).
(L) Representatives of DCX-expressing donor-derived $hN^+$ cells in the cerebral cortex at 4 months post-transplantation. Arrows indicate the bipolar processes of $DCX^+/hN^+$ cells. Scale bars, 20 μm.

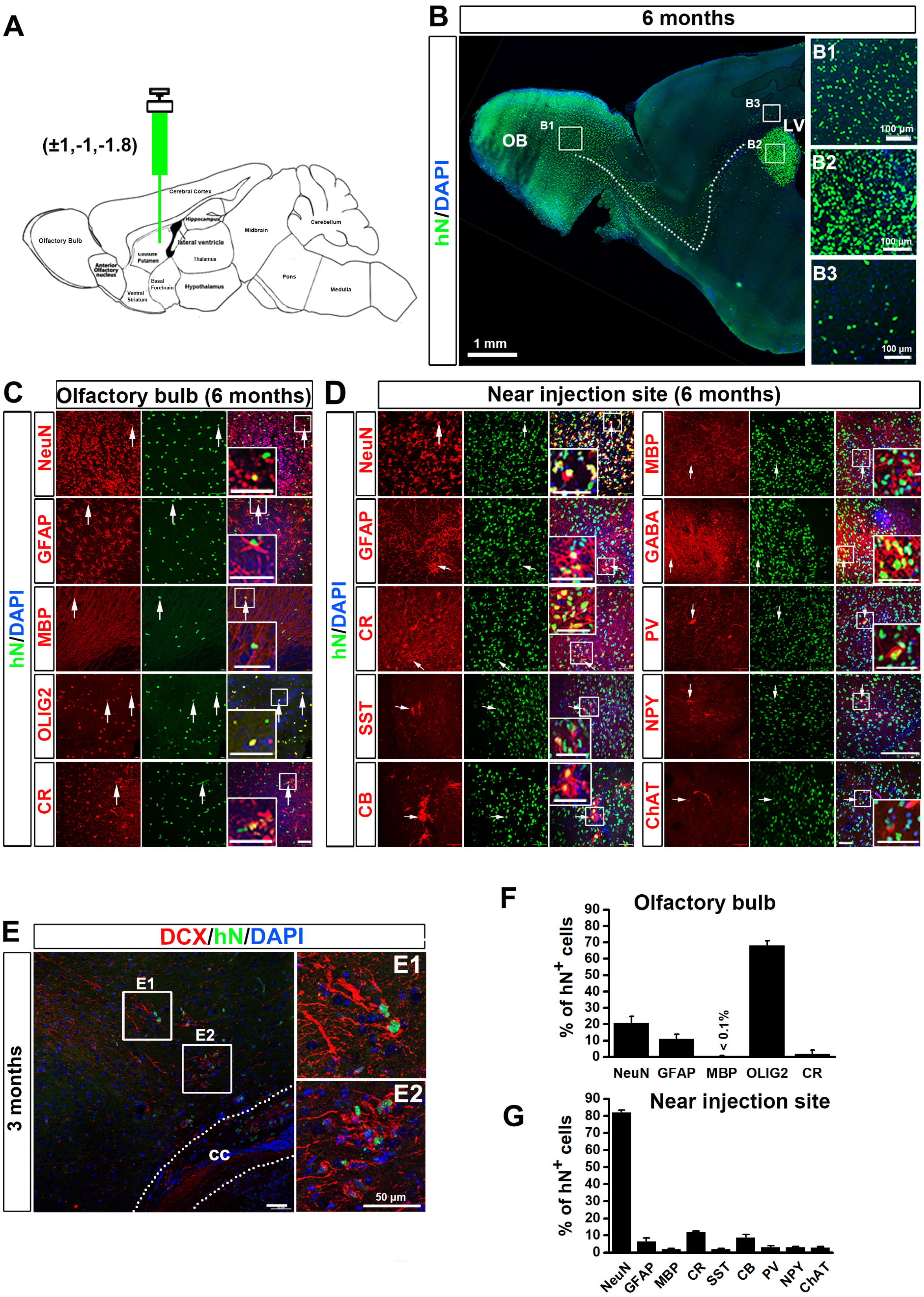

A
(±1,-1,-1.8)
Cerebral Cortex
Hippocampus
lateral ventricle
Midbrain
Cerebellum
Olfactory Bulb
Anterior Olfactory nucleus
Caudate Putamen
Thalamus
Basal Forebrain
Hypothalamus
Ventral Striatum
Pons
Medulla
B
6 months
hN/DAPI
OB
B1
B2
B3
LV
1 mm
100 µm
C
Olfactory bulb (6 months)
NeuN
GFAP
MBP
OLIG2
CR
D
Near injection site (6 months)
NeuN
GFAP
CR
SST
CB
MBP
GABA
PV
NPY
ChAT
E
DCX/hN/DAPI
3 months
E1
E2
cc
50 µm
F
Olfactory bulb
% of hN+ cells
< 0.1%
NeuN
GFAP
MBP
OLIG2
CR
G
Near injection site
% of hN+ cells
NeuN
GFAP
MBP
CR
SST
CB
PV
NPY
ChAT

**Figure S4. Related to Figure 4. Characterization of donor-derived cells after transplantation to the sites near subventricular zone**.
(A) A schematic diagram showing that control hiPSC-derived ventral forebrain NPCs are engrafted into the brains of P0 *rag*$1^{-/-}$ mice into the sites near the subventricular zone (SVZ).
(B) Representatives showing that donor-derived $hN^+$ cells have migrated along the rostral migratory stream to the olfactory bulb at 6 months after transplantation. Scale bar, 1 mm or 100 μm as indicated.
(C) Representatives of NeuN-, GFAP-, MBP-, OLIG2- and CR-expressing donor-derived $hN^+$ cells in the olfactory bulb at 6 months post-transplantation. Arrows indicate the donor-derived $hN^+$ cells expressing the indicated markers. Scale bars, 50 μm.
(D) Representatives of NeuN-, GFAP-, MBP-, CR-, SST-, CB-, PV-, NPY-, and ChAT-expressing donor-derived $hN^+$ cells in the regions near injection sites at 6 months post-transplantation. Arrows indicate the donor-derived $hN^+$ cells expressing the indicated markers. Scale bars, 50 μm.
(E) Representatives of DCX-expressing donor-derived $hN^+$ cells at 3 months post-transplantation in the cerebral cortex. CC, corpus callosum. Scale bars, 50 μm.
(F) Quantification of pooled data from control (Cont1 and Di-DS3) hiPSCs showing NeuN-, GFAP-, MBP-, OLIG2- and CR-expressing donor-derived $hN^+$ cells among the total $hN^+$ cells in the olfactory bulb at 6 months post-transplantation (n = 7 mice for each group). Data are presented as mean ± s.e.m.
(G) Quantification of pooled data from control (Cont1 and Di-DS3) hiPSCs showing NeuN-, GFAP-, MBP-, CR-, SST-, CB-, PV-, NPY-, and ChAT-expressing donor-derived $hN^+$ cells in the regions near injection sites among the total $hN^+$ cells at 6 months post-transplantation (n = 7 mice for each group). Data are presented as mean ± s.e.m.

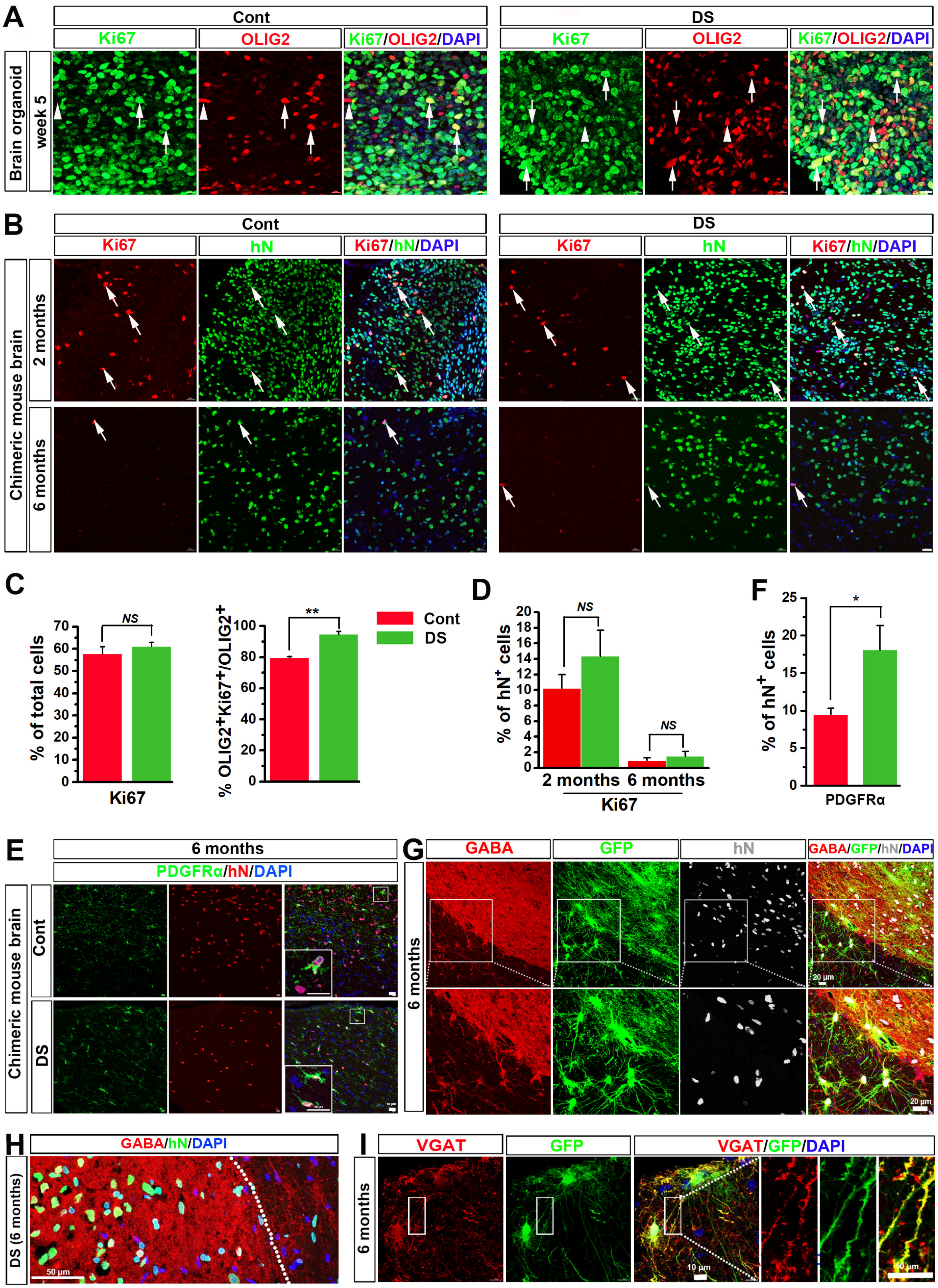
A
Cont
DS
Ki67
OLIG2
Ki67/OLIG2/DAPI
Brain organoid
week 5
B
Ki67
hN
Ki67/hN/DAPI
Chimeric mouse brain
2 months
6 months
C
% of total cells
NS
Ki67
% OLIG2+Ki67+/OLIG2+
**
Cont
DS
D
% of hN+ cells
NS
2 months
6 months
Ki67
F
*
PDGFRα
E
6 months
PDGFRα/hN/DAPI
Cont
DS
G
GABA
GFP
GABA/GFP/hN/DAPI
20 µm
H
GABA/hN/DAPI
DS (6 months)
50 µm
I
VGAT
VGAT/GFP/DAPI
10 µm

**Figure S5. Related to Figure 2 and Figure 4. Proliferation of the cells before and after transplantation and interneuron identity after transplantation.**

(A) Representatives of Ki67-expressing proliferating cells in control and DS 5-week-old ventral forebrain organoids. Arrows indicate OLIG2$^+$/Ki67$^+$ cells and arrowheads indicate OLIG2$^+$/Ki67$^-$ cells. Scale bars: 10 μm.

(B) Representatives of Ki67-expressing donor-derived hN$^+$ cells in the cerebral cortex at 2 and 6 months post-transplantation. Arrows indicate the donor-derived hN$^+$ cells expressing Ki67. The images were from confocal stitched tile scan. Scale bars, 20 μm.

(C) Quantification of pooled data from three pairs of control and DS hiPSCs showing the percentage of Ki67$^+$ cells in the total cells, as well as the percentage of Ki67$^+$/OLIG2$^+$ in total OLIG2$^+$ cells in 5-week-old control and DS ventral forebrain organoids (n = 4 and for each experiment, 4 to 6 organoids from each cell line are used). Student's *t* test. $^{**}P < 0.01$ and *NS* represents no significance. Data are presented as mean ± s.e.m

(D) Quantification of pooled data from control (Cont1 and Di-DS3) and DS (DS1 and TriDS3) hiPSCs showing the percentage of Ki67$^+$ cells among the total hN$^+$ cells in the cerebral cortex at 2 and 6 months post-transplantation (n = 7 mice for each group). Student's *t* test. *NS* represents no significance. Data are presented as mean ± s.e.m.

(E) Representatives of PDGFRα-expressing donor-derived hN$^+$ cells in the cerebral cortex at 6 months post-transplantation. Scale bars, 20 μm.

(F) Quantification of pooled data from control (Cont1 and Di-DS3) and DS (DS1 and Tri-DS3) hiPSCs showing the percentage of PDGFRα$^+$ cells among the total hN$^+$ cells in the cerebral cortex at 6 months post-transplantation (n = 7 mice for each group). Student's *t* test. $^*P < 0.05$. Data are presented as mean ± s.e.m.

(G) Representative images showing strong GABA staining in areas enriched of donor-derived hN$^+$ cells in the cerebral cortex at 6 months after transplantation. The cells are labeled by lenti-CMV-GFP, prior to transplantation. Scale bar, 20 μm.

(H) A representative image from DS chimeric mice showing high fluorescence intensity of GABA staining in areas enriched of donor-derived hN$^+$ cells in the cerebral cortex at 6 months after transplantation. Scale bar, 50 μm.

(I) Representative images showing GFP$^+$ donor-derived cells also express VGAT in the cerebral cortex at 6 months after transplantation. The cells are labeled by lenti-CMV-GFP, prior to transplantation. Scale bar, 10μm.

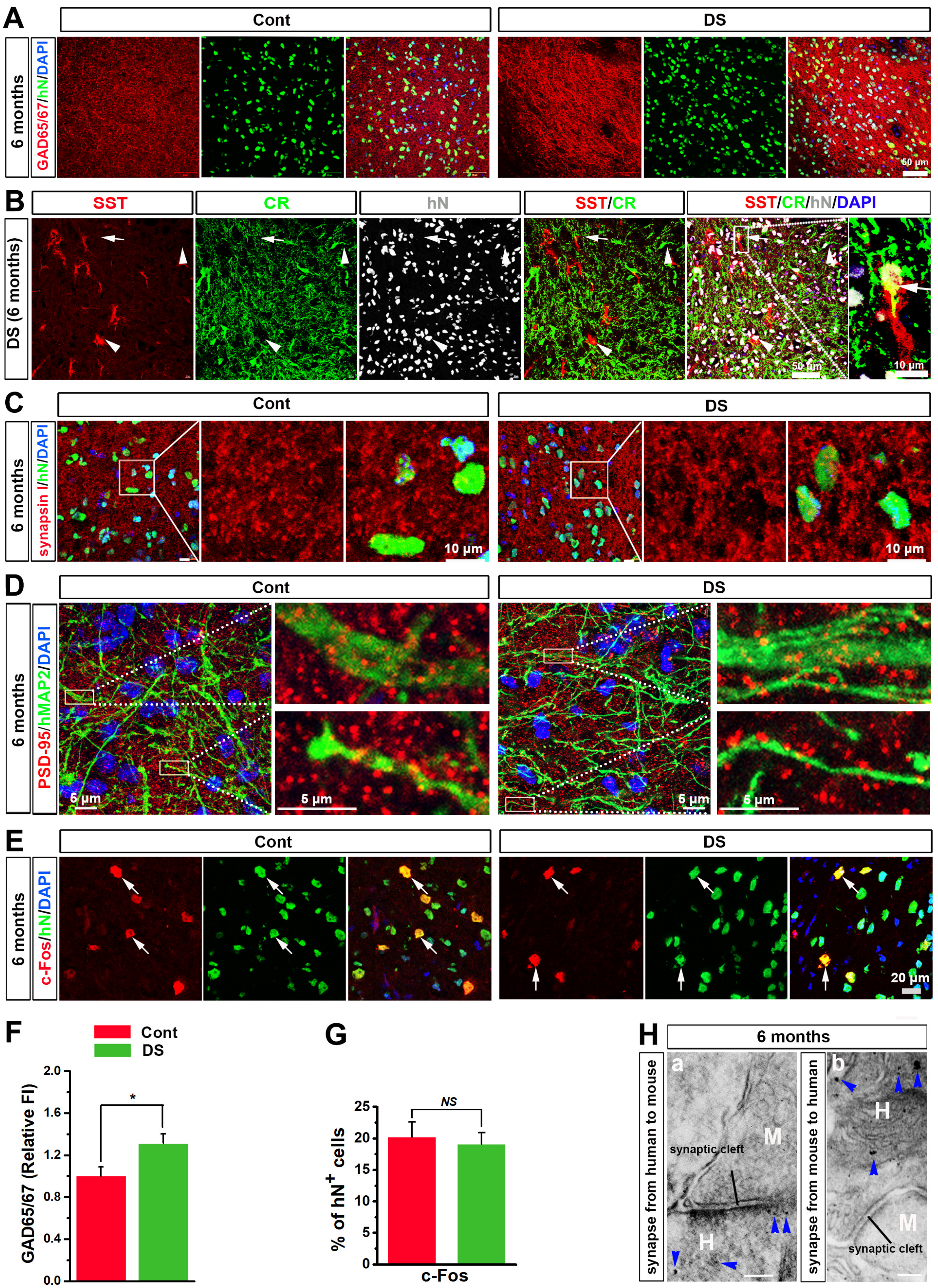
A
Cont
DS
6 months
GAD65/67/hN/DAPI
50 µm
B
SST
CR
hN
SST/CR
SST/CR/hN/DAPI
DS (6 months)
50 µm
10 µm
C
Cont
DS
6 months
synapsin I/hN/DAPI
10 µm
10 µm
D
Cont
DS
6 months
PSD-95/hMAP2/DAPI
5 µm
5 µm
5 µm
5 µm
E
Cont
DS
6 months
c-Fos/hN/DAPI
20 µm
F
Cont
DS
GAD65/67 (Relative FI)
2.0
1.6
1.2
0.8
0.4
0.0
*
G
% of hN+ cells
25
20
15
10
5
0
NS
c-Fos
H
6 months
synapse from human to mouse
a
M
synaptic cleft
H
synapse from mouse to human
b
H
M
synaptic cleft

**Figure S6. Related to Figure 4. Overabundance of GABAergic neurons in DS chimeric mouse brains and integration of donor cell-derived neurons into the brain.**
(A) Representatives of GAD65/67staining in the cerebral cortex at 6 months after transplantation of control or DS hiPSC-derived ventral forebrain NPCs. Scale bar, 50 μm.
(B) Representatives of SST, CR, and hN triple-staining in the cerebral cortex at 6 months after transplantation of DS hiPSC-derived ventral forebrain NPCs. Notably, few $hN^+$ human cells co-express SST and CR. Arrows indicate the donor-derived $hN^+$ cells co-expressing SST and CR. Arrowheads indicate the donor-derived $hN^+$ cells only expressing SST or CR. Scale bar, 50 μm or 10 μm in the original or enlarged images, respectively.
(C) Representatives of the $hN^+$ cell nuclei surrounded by the staining of presynaptic marker synapsin I in the cerebral cortex at 6 months after transplantation of control or DS hiPSC-derived ventral forebrain NPCs. Scale bar, 10 μm.
(D) Representatives of human-specific MAP2 (hMAP2)-expressing dendrites that co-localize with puncta of PSD-95 staining, a postsynaptic marker, in the cerebral cortex at 6 months after transplantation of control or DS hiPSC-derived ventral forebrain NPCs. Scale bar, 5 μm.
(E) Representatives of c-Fos-expressing donor-derived $hN^+$ cells in the cerebral cortex at 6 months post-transplantation of control or DS NPCs. Arrows show engrafted $hN^+$ cells expressing c-Fos. Scale bars, 20 μm.
(F) Quantification of pooled data from control (Cont1 and Di-DS3) and DS (DS1 and Tri-DS3) showing the fluorescence intensity (FI) of GAD65/67 staining in the cerebral cortex at 6 months after cell transplantation (n = 7 mice for each group). Student's *t* test. $^*P < 0.05$. Data are presented as mean ± s.e.m.
(G) Quantification of pooled data from control (Cont1 and Di-DS3) and DS (DS1 and Tri-DS3) hiPSCs showing the percentage of c-$Fos^+$ cells among the total $hN^+$ cells in the cerebral cortex at 6 months post-transplantation (n = 7 mice for each group). Student's *t* test. *NS* represents no significance. Data are presented as mean ± s.e.m.
(H) Representative electron microscopy images of synaptic terminals formed between human neurons labeled by DAB staining against a human-specific marker STEM121 and mouse neurons that were not labeled by the DAB staining. (a) synaptic contacts from human transplant to mouse host tissue and (b) synapse from mouse neurons to human neurons at 6 months post-transplantation. Synaptic cleft structure was seen in the synaptic terminals. "H" denotes human neurons, "M" denotes mouse neurons, and arrowheads indicate DAB electron-dense precipitates. Scale bar represents 100 nm.

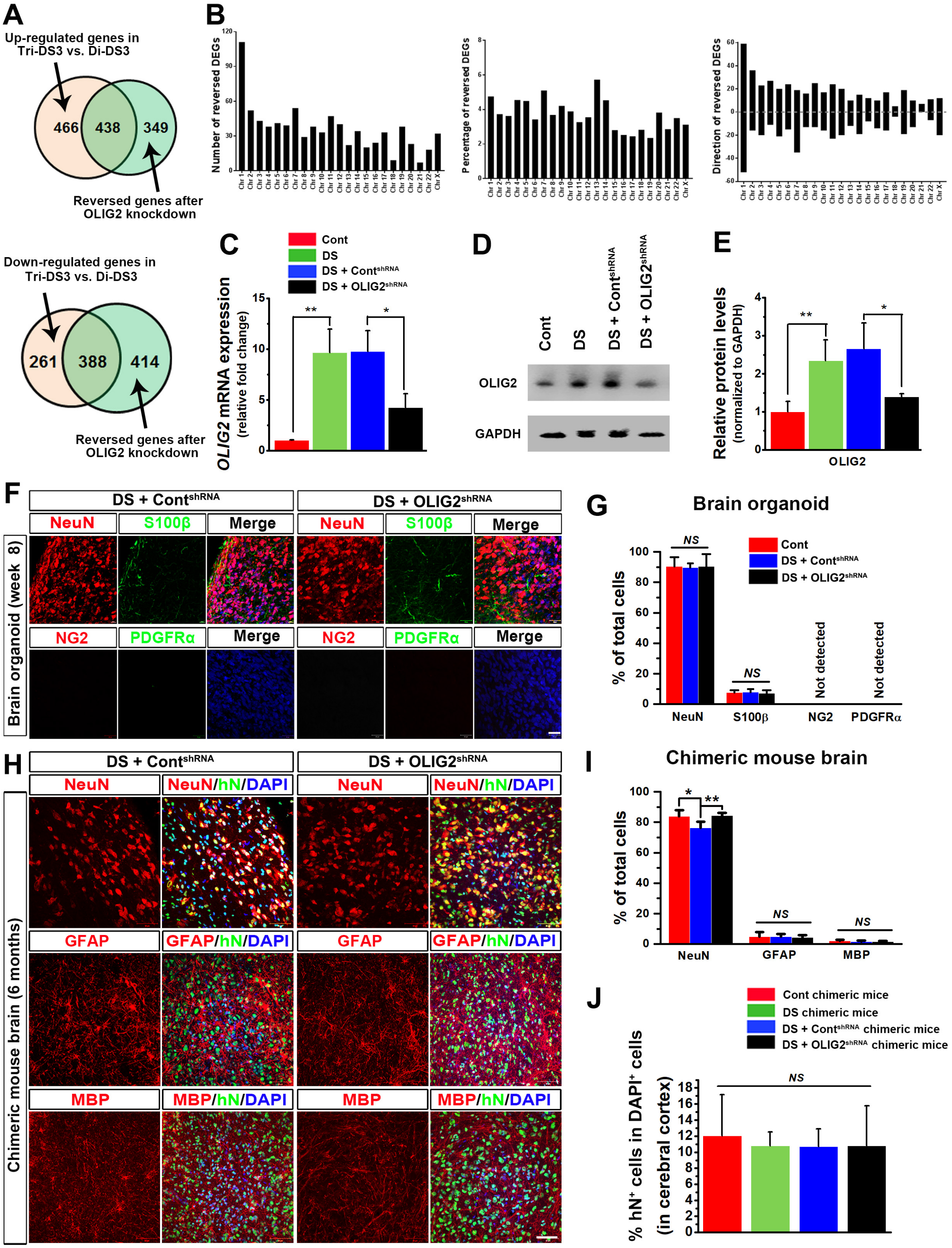
A
Up-regulated genes in Tri-DS3 vs. Di-DS3
466
438
349
Reversed genes after OLIG2 knockdown
Down-regulated genes in Tri-DS3 vs. Di-DS3
261
388
414
Reversed genes after OLIG2 knockdown
B
Number of reversed DEGs
Percentage of reversed DEGs
Direction of reversed DEGs
C
Cont
DS
DS + Cont$^{shRNA}$
DS + OLIG2$^{shRNA}$
OLIG2 mRNA expression (relative fold change)
D
Cont
DS
DS + Cont$^{shRNA}$
DS + OLIG2$^{shRNA}$
OLIG2
GAPDH
E
Relative protein levels (normalized to GAPDH)
OLIG2
F
DS + Cont$^{shRNA}$
DS + OLIG2$^{shRNA}$
NeuN
S100β
Merge
NG2
PDGFRα
Brain organoid (week 8)
G
Brain organoid
% of total cells
NS
Not detected
NeuN
S100β
NG2
PDGFRα
H
NeuN/hN/DAPI
GFAP
GFAP/hN/DAPI
MBP
MBP/hN/DAPI
Chimeric mouse brain (6 months)
I
Chimeric mouse brain
% of total cells
NeuN
GFAP
MBP
J
Cont chimeric mice
DS chimeric mice
DS + Cont$^{shRNA}$ chimeric mice
DS + OLIG2$^{shRNA}$ chimeric mice
% hN$^+$ cells in DAPI$^+$ cells (in cerebral cortex)
NS

**Figure S7. Related to Figure 5 and Figure 6. Reversed expression of differentially expressed genes (DEGs) in Tri-DS3 organoids by OLIG2 knockdown and differentiation of DS ventral forebrain NPCs after OLIG2 knockdown in the organoids and chimeric mouse brains.**

(A) Venn diagrams showing 438 upregulated and 388 downregulated DEGs are reversed after *OLIG2* knockdown (KD) in 5-week-old Di-DS3 and Tri-DS3 organoids.

(B) The total number of reversed DEGs, percentage of reversed DEGs, and number of up-regulated or down-regulated of DEGs that are reversed after OLIG2 knockdown on each chromosome in 5-week-old Di-DS3 and Tri-DS3 organoids.

(C) qPCR analysis of *OLIG2* mRNA expression in 5-week-old control, DS, DS + $Cont^{shRNA}$ and DS + $OLIG2^{shRNA}$ organoids. The data are pooled from control (Cont1 and Di-DS3) hiPSCs, DS (DS1 and Tri-DS3) hiPSCs, and the two lines of DS hiPSCs infected with lentiviruses that carry control shRNA or OLIG2 shRNA. The experiments are repeated for three times (n = 3) and for each experiment, 20 to 30 organoids from each line are used. Student's *t* test. $^{**}P < 0.01$, $^{*}P < 0.05$. Data are presented as mean ± s.e.m.

(D) Western blotting analysis of OLIG2 expression in 5-week-old control, DS, DS + $Cont^{shRNA}$ and DS + $OLIG2^{shRNA}$ organoids.

(E) Quantification data showing OLIG2 expression by western blotn in 5-week-old control, DS, DS + $Cont^{shRNA}$ and DS + $OLIG2^{shRNA}$ organoids. The data are pooled from control (Cont1 and Di-DS3) hiPSCs, DS (DS1 and Tri-DS3) hiPSCs, and the two lines of DS hiPSCs infected with lentiviruses that carry control shRNA or OLIG2 shRNA. The experiments are repeated for three times (n = 3) and for each experiment, 30 to 40 organoids from each line are used. Student's *t* test. $^{**}P < 0.01$, $^{*}P < 0.05$. Data are presented as mean ± s.e.m.

(F) Representatives of NeuN-, S100β-, NG2-, and PDGFRα-expressing cells in 8-week-old DS + $Cont^{shRNA}$ and DS + $OLIG2^{shRNA}$ organoids. Scale bars: 20 μm.

(G) Representatives of NeuN-, GFAP-, and MBP-expressing donor-derived $hN^+$ cells in the cerebral cortex at 6 months after transplantation of DS + $Cont^{shRNA}$ and DS + $OLIG2^{shRNA}$ ventral forebrain NPCs. Scale bars, 50 μm.

(H) Quantification of percentage of $NeuN^+$, $S100\beta^+$, $NG2^+$, and $PDGFR\alpha^+$ in 8-week-old control (Cont), DS + $Cont^{shRNA}$, and DS + $OLIG2^{shRNA}$ organoids. The data are pooled from control (Cont1 and Di-DS3) hiPSCs, DS (DS1 and Tri-DS3) hiPSCs, and the two lines of DS hiPSCs infected with lentiviruses that carry control shRNA or OLIG2 shRNA. The experiments are repeated for three times (n = 3) and for each experiment, 8 to10 organoids from each line are used. Data are presented as mean ± s.e.m.

(I) Quantification of the percentage of $NeuN^+$, $GFAP^+$, and $MBP^+$ expressing cells among total $hN^+$ cells in the cerebral cortex at 6 months after transplantation of control (Cont), DS + $Cont^{shRNA}$, or DS + $OLIG2^{shRNA}$ ventral forebrain NPCs (n = 7 mice for each group). The data are pooled from mice received transplantation of the NPCs derived from control (Cont1 and Di-DS3) hiPSCs, DS (DS1 and Tri-DS3) hiPSCs, and the two lines of DS hiPSCs infected with lentiviruses that carry control shRNA or OLIG2 shRNA. One-way ANOVA test, $^{*}P < 0.05$, $^{**}P < 0.01$, and *NS* represents no significance. Data are presented as mean ± s.e.m.

(J) Quantification of the percentage of $hN^+$ cells in total $DAPI^+$ cells in the cerebral cortex at 6 months post-transplantation in control (Cont), DS + $Cont^{shRNA}$, or DS + $OLIG2^{shRNA}$ chimeric mouse groups (n = 7-8 mice for each group). The data are pooled from mice received transplantation of the NPCs derived from control (Cont1 and Di-DS3) hiPSCs, DS (DS1 and Tri-DS3) hiPSCs, and the two lines of DS hiPSCs infected with lentiviruses that carry control shRNA or OLIG2 shRNA. One-way ANOVA test. *NS* represents no significance. Data are presented as mean ± s.e.m.

**Table S1. Related to Figure 2. Expression of Olig1 and 2 and major physiological phenotypes in iTg-Nes transgenic mice with mis-expression of Olig2 and mouse genetic models of DS.**

| Mouse models | Genotype and phenotypic expression of *Olig* genes | Changes in the number of GABAergic neurons | Synaptic plasticity | Behavioral phenotypes |
|---|---|---|---|---|
| **iTg-Nes** | Olig2 is overexpressed and mis-expressed in cortical neural stem cells (Liu et al., 2015) | Increased but observed abnormally massive neuronal cell death (Liu et al., 2015) | Not examined | Locomotion abnormalities (Liu et al., 2015) |
| **Ts65Dn** | Trisomic for *Olig1* and *2;* Olig1 and 2 are overexpressed (Aziz et al., 2018; Chakrabarti et al., 2010) | Increased (Chakrabarti et al., 2010; Hernandez-Gonzalez et al., 2015; Perez-Cremades et al., 2010) | Deficient long-term potentiation (LTP) in CA1 and DG (Costa and Grybko, 2005; Fernandez et al., 2007; Garcia-Cerro et al., 2014; Kleschevnikov et al., 2012; Kleschevnikov et al., 2004; Lysenko et al., 2014; Siarey et al., 1997) | Locomotion abnormalities and Learning and memory deficits (Braudeau et al., 2011; Escorihuela et al., 1995; Escorihuela et al., 1998; Faizi et al., 2011; Hyde et al., 2001; Kleschevnikov et al., 2012; Lysenko et al., 2014; Olson et al., 2007; Rueda et al., 2010; Smith et al., 2014; Stasko and Costa, 2004) |
| **Ts1Cje** | Trisomic for *Olig1* and *2;* Olig2 expression is normal (may be related to chromatin state) (Aziz et al., 2018) | Not examined, but observed Increase GAD65$^+$ inhibitory input in the hippocampus (Belichenko et al., 2007). | Deficient LTP in CA1 and DG (Belichenko et al., 2007; Siarey et al., 2005) | Learning and memory deficits (less severe than Ts65Dn)(Belichenko et al., 2007; Fernandez and Garner, 2007; Sago et al., 1998; Sago et al., 2000) |
| **Tc1** | Trisomic for *Olig1* and *2;* expression of Olig1 and 2 is not examined | Not examined | Deficient LTP in DG (Morice et al., 2008; O'Doherty et al., 2005) | Locomotion abnormalities and mild learning impairment(Galante et al., 2009; Morice et al., 2008; O'Doherty et al., 2005) |
| **TTS** | Trisomic for *Olig1* and *2;* expression of Olig1 and 2 is not examined | Not examined | Deficient LTP in CA1 and DG (Belichenko et al., 2015; Yu et al., 2010a) | Locomotion abnormalities and Learning and memory deficits (Belichenko et al., 2015) |
| **Dp(16)1Yey** | Trisomic for *Olig1* and *2,* but Olig2 expression is normal (may be related to chromatin state)(Olmos-Serrano et al., 2016) | Decreased (Olmos-Serrano et al., 2016) | Deficient LTP in CA1 (Yu et al., 2010b) | Locomotion abnormalities and Learning and memory deficits (Olmos-Serrano et al., 2016; Yu et al., 2010b) |
| **Dp(10)1Yey** | Disomic for *Olig1* and *2* | Not examined | No change (Yu et al., 2010b) | No change(Yu et al., 2010b) |
| **Dp(17)1Yey** | Disomic for *Olig1* and *2* | Not examined | Increased LTP in CA1(Yu et al., 2010b) | No Change(Yu et al., 2010b) |
| **Ts1Rhr** | Disomic for *Olig1* and *2* | Not examined | Deficient LTP in fascia dentata (Belichenko et al., 2009; Olson et al., 2007) | Locomotion abnormalities and learning impairment (less severe than Ts65Dn)(Belichenko et al., 2009) |
| **Ts1Yah** | Disomic for *Olig1* and *2* | Not examined | Increased LTP in CA1(Pereira et al., 2009) | Defects in working memory; Improvement in spatial memory(Pereira et al., 2009) |

**Table S2. Related to Figure 2. Basic information of Down syndrome patient fibroblasts from Coriell Institute for Medical Research.**

| Coriell's cat. no. | Sex | Age | Race | Cell type | Karyotype | Generated iPSC lines |
|---|---|---|---|---|---|---|
| GM04616 | Female | 3 Days | Caucasian | Fibroblast | 47,XX,+21 | DS1 iPSCs |
| AG08942 | Male | 21 Years | Caucasian | Fibroblast | 47,XY,+21 | DS2 iPSCs |
| AG06872 | Female | 1 Year | Caucasian | Fibroblast | 47,XX,+21 | Isogenic Di-DS and Tri-DS iPSCs |

**Table S3. Related to Figure 3. Basic information of the human brain tissues obtained from the University of Maryland Brain and Tissue Bank (UMB).**

| UMB no. | Disorder | Sex | Age (Days) | Cell type | Race | Post mortem interval (hours) |
|---|---|---|---|---|---|---|
| **832** | Trisomy 21 | Male | 339 | Cerebral cortex | African American | 23 |
| **1282** | Trisomy 21 | Female | 186 | Cerebral cortex | Caucasian | 28 |
| **4783** | Trisomy 21 | Female | 261 | Cerebral cortex | Caucasian | 37 |
| **M1931M** | Control | Male | 332 | Cerebral cortex | Caucasian | 10 |
| **4361** | Control | Female | 236 | Cerebral cortex | African American | 27 |
| **1443** | Control | Female | 332 | Cerebral cortex | African American | 10 |

**Table S4. Related to Figure 5. Taqman Primers.**

| Gene | Gene expression assay catalog number |
|---|---|
| *OLIG2* | Hs00300164_s1 |
| *OLIG1* | Hs00744293_s1 |
| *DLX1* | Hs00698288_m1 |
| *DLX2* | Hs00269993_m1 |
| *DLX5* | Hs00193291_m1 |
| *DLX6* | Hs00231999_m1 |
| *LHX6* | Hs01030943_m1 |
| *LHX8* | Hs00418293_m1 |
| *SOX6* | Hs00264525_m1 |
| *GAPDH* | Hs02758991_g1 |

**Table S5. Related to Figure 5. ChIP-PCR Primers.**

| Human LHX promotor | Forward | Reverse |
|---|---|---|
| **-162 to -345** | 5'- CCTCTGCTACTGCACACAAT-3' | 5'- ATCTGCAAGGAAAAGGAGAGA-3' |
| **-479 to -696** | 5'- GCTTGAAAGATGCAGGAACT-3' | 5'- ATCCAGGAAGCTTGTAGAGC-3' |
| **-860 to -1643** | 5'- CTTAGGCACTCAGAGTAGGT-3' | 5'- AGCCCTCCGTGTCGGATGTC-3' |
| **-1149 to -1321** | 5'- TTCTAGGCCAGAGAGGAT-3' | 5'- GTCCAGAGTCTGTCTATGGG-3' |
| **-1384 to -1521** | 5'- GACAAGGGCCTGTTTCTGCT-3' | 5'- ATGAAGACTCCATCTACTG-3' |
| **-1557 to -1701** | 5'- GCTGAGTTCGCAAATCTTCT-3' | 5'- GCTGGTAAAACCATTCCAGT-3' |
| **-1756 to -1890** | 5'- TTAGCCACATCAGGGTGTTAT-3' | 5'- GGTATTTGAGGGAAATAAATGGG-3' |
| **Human DLX1 promotor** | **Forward** | **Reverse** |
| **-25 to -186** | 5'- CGCTCCAGAGGCAGAGGT -3' | 5'- AGCCGCTGGTTGGTTCCTG -3' |
| **-328 to -452** | 5'- GCACTGCTTTAATCAGACC -3' | 5'- GAGTGAGTGAGTTCGCAGG -3' |
| **-642 to -830** | 5'- AGGTGAGGAAGATCCTGGGT -3' | 5'- AGATCTGAGCCCCTAG -3' |
| **-1126 to -1251** | 5'- CATCAGTGTTATGCTAATGG -3' | 5'- ACAAGATCATTTCCCCGTT-3' |
| **-1467 to -1621** | 5'- TTGCTCCAACCACTCTGCCT -3' | 5'- AACGACAGCACCGATGTAAT -3' |
| **-1670 to -1921** | 5'- AGGGCACGCGCGAGAGACCT -3' | 5'- AGCGGCTCCCGGAGTTGCCTG -3' |

**Table S6. Related to Figure 5. Primers for luciferase reporter assay.**

| | Forward | Reverse |
|---|---|---|
| **LHX8 (-1 to -2000)** | 5'-GTAGCTAGCCCTTAAAAAGGCATCGTATG -3' | 5'-GTAAGATCTGCCCACTTCCGCTGAGCAGC-3' |
| **DLX1 (-10 to -1250)** | 5'- GTAGCTAGCATCAGTGTTATGCTAATGG -3' | 5'- TACAAGCTTGCCTCCCTCGCGCGCTTTCC -3' |
